# Iron Oxide Nanoparticles Coated with Biodegradable Block-Copolymer PDMAEMA-b-PMPC and Functionalized with Aptamer for HER2 Breast Cancer Cell Identification

**DOI:** 10.1101/2023.06.06.543859

**Authors:** Cyro von Zuben de Valega Negrão, Natália Neto Pereira Cerize, Amauri da Silva Justo-Junior, Raquel Bester Liszbinski, Giovanna Pastore Meneguetti, Larissa Araujo, Silvana Aparecida Rocco, Kaliandra de Almeida Gonçalves, Daniel Reinaldo Cornejo, Patrícia Leo, Caio Perecin, Douglas Adamoski, Sandra Martha Gomes Dias

## Abstract

Hybrid nanoparticles have shown promise in biomedical applications; however, their seamless integration into clinical settings remains challenging. Here, we introduce a novel metal oxide polymer hybrid nanoparticle (NP) with a high affinity for nucleic acids. Iron oxide nanoparticles (IONP) were initially synthesized via the co-precipitation method and subjected to comprehensive characterization. Subsequently, block copolymers were synthesized using the Reversible Addition−Fragmentation Chain Transfer (RAFT) technique, employing the zwitterionic PMPC (Poly (2 Methacryloyloxyethyl Phosphorylcholine)) and the cationic PDMAEMA (Poly(2 (Dimethylamino) Ethyl Methacrylate)) with varying degrees of polymerization. In vitro cytotoxicity studies demonstrated the biocompatibility of the synthesized nanoparticles, with no observed toxicity up to a concentration of 150 µg/mL. The cationic polymer PDMAEMA facilitated the facile coating of IONP, forming the IONPP complex, consisting of a 13.27 metal core and a 3.1 nm block-copolymer coating. Subsequently, the IONPP complex was functionalized with a DNA aptamer specifically targeting the human epidermal growth factor receptor 2 (HER2) in breast cancer, forming IONPPP. The block-copolymer exhibited an EC_50_ of 7.07 µg/mL and demonstrated enhanced recognition efficiency in HER2-amplified SKBR3 cells. Our study presents a comprehensive IONPPP characterization capable of binding short DNA sequences and targeting proteins such as HER2. This newly developed nanoparticle holds significant potential for cancer cell identification and isolation, offering promising prospects in cancer research and clinical applications.

**Statement of significance:** Despite recent advancements in biomedical research, developing sensitive and specific tools for recognizing biological motifs, such as cell receptors and proteins in complex biological solutions, remains a challenge. Furthermore, current approaches often rely on complex biological derivatives like antibodies, lacking a cost-effective delivery strategy. Our study proposes creating and characterizing a novel hybrid metal oxide polymer nanoparticle named IONPPP, functionalized with a DNA aptamer designed to recognize HER2-positive cells. HER2 is a clinically actionable marker for gastric, gastroesophageal, and, particularly, breast cancers. This unique combination of a metal core with an external polymeric structure offers the potential for identification, isolation, and even theragnostic applications, benefiting from its low toxicity and high specificity.

**Graphical Abstract:** 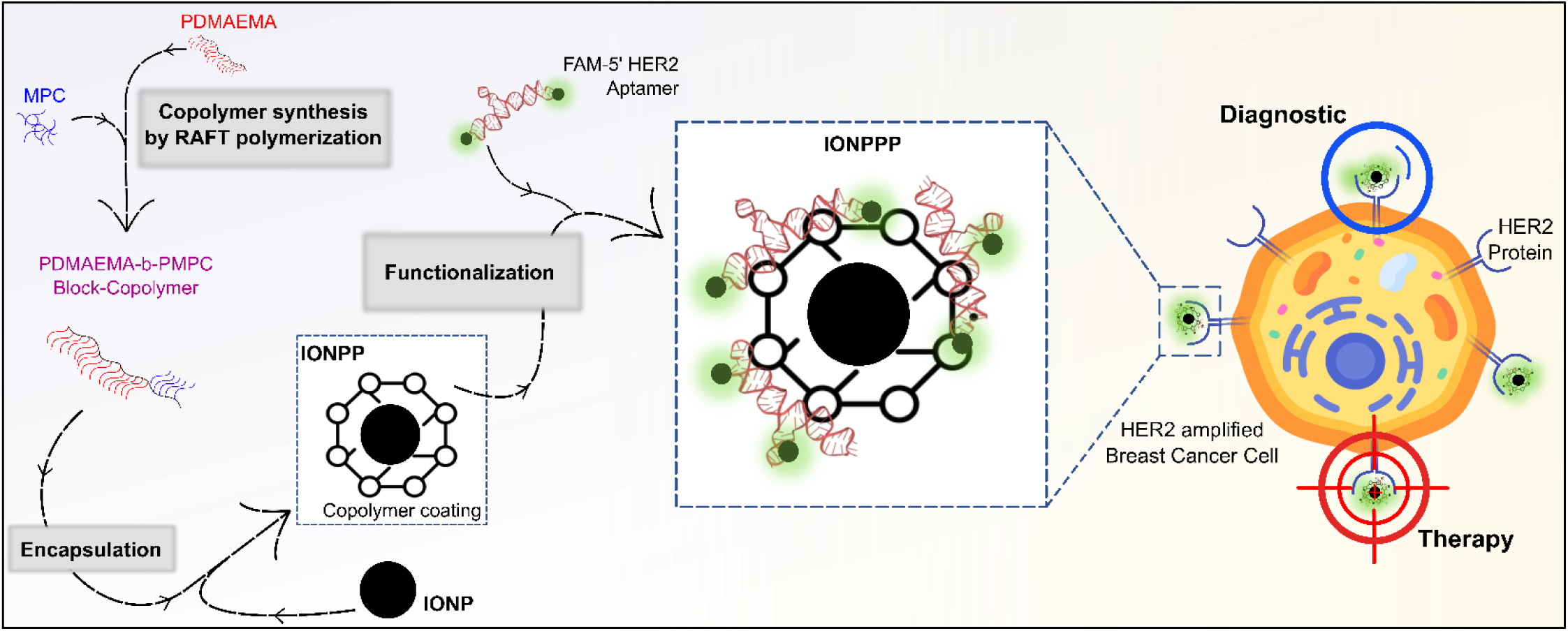

## 3. Introduction

Iron oxide nanoparticles (IONPs) are nanostructured assemblies of iron and oxygen atoms with remarkable properties, including superparamagnetism (Ajinkya et al., 2020; Ali et al., 2016). Over the past few decades, IONPs have found applications in various scientific fields, particularly in biomedical research (Gupta & Gupta, 2005; W. Wu et al., 2016). Notably, they have been extensively utilized in magnetic hyperthermia for therapeutic purposes and magnetic resonance imaging (MRI) for diagnostic imaging (Israel et al., 2020; Nosrati et al., 2019; Thakor et al., 2016). Additionally, owing to their ability to serve as drug carriers and promote drug release, IONPs have gained attention in the field of controlled drug delivery (da Silva et al., 2018; Norouzi et al., 2020). Furthermore, leveraging their magnetic properties, IONPs have emerged as a promising tool for diagnosing cancer cells (Onishi et al., 2022; Su et al., 2020; Zhao et al., 2020).

Despite the recent advancements in the synthesis and purification methods of IONPs, several challenges still need to be addressed. Firstly, IONPs exhibit a short circulation half-life, limiting their efficacy in various biomedical applications. Secondly, issues related to large-scale scalability and reproducibility hinder their widespread use. Additionally, IONPs tend to aggregate when uncoated, compromising their stability and performance (W. Wu et al., 2008, 2016). To overcome these challenges, the encapsulation of IONPs with other molecules, such as biodegradable polymers, has emerged as a promising approach (Sperling & Parak, 2010). In addition, the polymeric coating promotes improved colloidal stability, preventing aggregation and ensuring long-term stability. Furthermore, combining metal and polymer components allows for developing theragnostic nanocomposites, integrating diagnostic and therapeutic functionalities. Lastly, the hybrid structure enhances the biocompatibility of the nanoparticles, expanding their potential applications (K. Wu et al., 2019; Zhao et al., 2020).

Among different strategies for polymer synthesis, reversible addition−fragmentation chain transfer (RAFT) has demonstrated several advantages, such as being a well-controlled, versatile, and free-of-toxic element polymerization process (Alidedeoglu et al., 2009; Keddie, 2014; Perrier, 2017). Poly (2 methacryloyloxyethyl phosphorylcholine) (PMPC) is a zwitterionic polymer known for its high biocompatibility and low immunogenicity. In some studies, PMPC has shown an improved toxicity profile compared to poly (ethylene glycol) (PEG) (Auriemma et al., 2021; Goda et al., 2015a; Luongo et al., 2017; Thi et al., 2020). Poly(2-(dimethylamino) ethyl methacrylate) (PDMAEMA) is a cationic hydrophilic polymer commonly used as a carrier for gene and drug delivery systems (Gautier et al., 2013; Tzankova et al., 2016). The block-copolymer PDMAEMA-b-PMPC represents a synergistic combination that harnesses the unique properties of both polymers. PMPC provides protection, biocompatibility, and structural conformation, while PDMAEMA offers flexibility and the ability to selectively bind and protect nucleic acids from chemical and biological degradation. Furthermore, the PDMAEMA component enables tight retention of nucleic acids in non-acidic biological environments, such as the bloodstream, and controlled release within specific organelles like the lysosome due to the proton effect (Gautier et al., 2013; Tzankova et al., 2016). This combination of properties makes PDMAEMA-b PMPC an excellent candidate for enhancing the stability, biocompatibility, and efficient delivery of nucleic acids in various biomedical applications. Using this novel complex for functionalizing IONP particles opens opportunities for creating multifunctional nanocomposites capable of encapsulating 3D-structured nucleic acids with biological effects spanning diagnostics to therapeutics.

Aptamers are artificial single-stranded RNA or DNA sequences consisting of dozens of base pairs with biochemical functions as standalone molecules (Xiao & Farokhzad, 2012). These molecules have been utilized in various applications since the proposition of in vitro evolution to improve and select functional molecules as self replicating nucleic acids (Mills et al., 1967). To design and select functional sequences suitable for aptamer development, the Systematic Evolution of Ligands by Exponential Enrichment (SELEX) method was established as a highly efficient and practical tool (Sheikh et al., 2022). This tool selects the most appropriate sequence with high affinity and selectivity to target since it can fold into tertiary structures that bind precisely to proteins through salt bridges, electrostatic interaction, Van der Waals forces, hydrogen bonding, and aromatic ring stacking (Sheikh et al., 2022). Furthermore, aptamers are non immuno-responsive, chemically versatile, highly stable in different pHs, and cost effective compared to antibodies (Fu & Xiang, 2020; Xiao & Farokhzad, 2012). Their 10-30 kDa size and negative charge also make them ideal for nanoparticle complexing. The resulting NP-aptamer complexes can be used to specifically bind to various targets, including macromolecules and their carriers, as cancer cells, holding great promise in the fields of medicine and cancer research (Fu & Xiang, 2020; M. Liu et al., 2022; Sheikh et al., 2022; Xiao & Farokhzad, 2012). Aptamers have demonstrated their potentials in both diagnostics and therapy, such as binding to circulating tumor cells for early metastasis detection and blocking the activity of specific membrane proteins to induce effects like apoptosis and disruption of cancer cell growth and division. Clinical trials, including the phase II trial of AS1411, highlight the progress and potential of aptamer-based therapies (Bates et al., 2009; Hassan et al., 2016).

The tyrosine kinase HER2, which stands for human epidermal growth factor receptor 2 (*ERBB2* human gene), is an orphan receptor that plays a crucial role in cell growth and division. Breast cancer patients exhibiting high levels of HER2 expression often present with more aggressive and invasive tumors, leading to increased disease progression and reduced overall survival rates (Boulbes et al., 2015; Varty et al., 2021). In addition, overexpression or amplification of the HER2 gene leads to excessive production of HER2 protein, which can promote uncontrolled cell proliferation and tumor growth (Boulbes et al., 2015). Therefore, accurate identification and targeting of HER2 positive cells are essential for improving breast cancer treatment strategies and patient outcomes(Boulbes et al., 2015; Varty et al., 2021). However, there is an urgent need to create new alternatives to diagnose HER2^+^ cells in the clinical once testing has had high error rates (Gutierrez & Schiff, 2011).

Here, we present a new multifunctional nanoparticle composed of an iron oxide nanoparticle coated with a block-copolymer PDMAEMA-b-PMPC (IONPP) and functionalized with an anti-HER2 DNA aptamer (IONPPP). (Jiang et al., 2017; Liang et al., 2018; Poturnayová et al., 2019; Saleh et al., 2019; Vajhadin et al., 2022; K. Wang et al., 2015). Our innovative strategy leverages the synergistic properties of PMPC and PDMAEMA. PMPC enhances nanoparticle (NP) biocompatibility and stability in colloidal solutions, making them suitable for biological applications. On the other hand, PDMAEMA facilitates interaction with nucleic acid molecules, providing a conducive environment for aptamer binding. Our goal is to augment the efficacy of nucleic acid aptamers, not just in terms of binding efficiency but also in their targeted delivery to tissues. This nanostructure optimizes therapeutic applications, combining stability, biocompatibility, and effective nucleic acid delivery. Finally, the metal nanocomponent was essential to increase the sensitivity to capture HER2^+^ tumor cells. Thus, this multifunctional hybrid nanoparticle showcased in our study shows significant potential as a diagnostic tool in cancer diagnosis by integrating precise targeting and efficient cell capture.

## 4. Materials and Methods

### 4.1. Materials

Iron (III) chloride hexahydrate (FeCl_3_·6H_2_O, ≥ 98%), Iron(II) sulfate heptahydrate (Fe_2_SO_4_.7H_2_O, ≥ 98%), sodium sulfite (Na_2_SO_3_, ≥ 98%), sodium hydroxide (NaOH, ≥ 98%), 2-methacryloyloxyethyl phosphorylcholine (MPC, 97%,), 2-(Dimethylamino) Ethyl Methacrylate (DMAEMA, 98%), 4-cyano-4-(phenylcarbonothioylthio)pentanoic acid (CPA, 97%), 4,4′-azobis(cyanovaleric acid) (ACVA, ≥ 98%), Methanol-d (CD3OD, ≥ 99.8 atom %D, contains 0.03 % (v/v) TMS, ethyl alcohol (CH_3_CH_2_OH, ≥ 99%), hydrochloric acid (HCl, ≥ 99%), methanol (CH_3_OH, ≥ 99%), were purchased from Sigma-Aldrich (Merck). HER2 aptamer was synthesized by the LQS (Synthetic Chemistry Laboratory) at LNBio. All chemicals were used without further purification, and ultrapure water (18 MΩ·cm) was employed to prepare solutions and wash. For the biological characterization and application, we used propidium iodide (Sigma-Aldrich), RPMI 1640 medium (Sigma-Aldrich), 3-(4,5-Dimethyl-2-thiazolyl)-2,5-diphenyl-2H tetrazolium Bromide – MTT (Sigma-Aldrich), MitoTracker probe (Life Technologies), and DAPI (Sigma-Aldrich).

### 4.2. Magnetic nanoparticle synthesis

The magnetite synthesis was performed by the classical method co-precipitation (Massart, 1981) by mixing FeCl_3_.6H_2_O (0.032M, 10 mL) and Fe_2_SO_4_.7H_2_O (0.018M, 10 mL) salts in 30 mL of deionized H_2_O, producing a red-colored [Fe-SO_3_] complex. After, NaOH (0.5M, 10mL) was quickly added to the aqueous solution by syringe, changing the solution color to black. After 30 min under vigorous agitation at room temperature, the product was separated by centrifugation (10.000 RPM, 15 min) and washed with water five times to obtain the suspension of iron oxide nanoparticles (IONP). All steps were degassed with argon for 20 min.

### 4.3. RAFT Polymerization of (PMPC and PDMAEMA)

The amphiphilic and cationic polymers were synthesized by RAFT polymerization. PMPC was obtained by first adding 4-cyano-4-(phenylcarbonothioylthio)pentanoic acid (CPA) and MPC in acetate buffer (3 mM, pH 5.4) to a round bottom flask. Separately, 4,4′-azobis(cyanovaleric acid (ACVA) was diluted in ethanol 100%. After mixing MPC/ACVA and ACVA, the solution was degassed with argon for 20 min. The reaction occurred at 65°C for 20 h under magnetic stirring. For PDMAEMA synthesis, DMAEMA, CPA, and ACVA were diluted in methanol and the reaction occurred at 60°C. The PMPC purification was performed by solvent evaporation under air and washing the polymer with acetone (40 mL) at 5000 rpm for 10 min three times. Specifically, for PDMAEMA, dialysis was performed using 3.5-KDa membrane at 4°C overnight. Finally, both polymer products were dried overnight in a vacuum oven at 40°C. ^1^H NMR and FT IR confirmed the synthesis. The reagent amount for all polymer syntheses is in Table 1.

**Table 1.** Reagents used to synthesize the zwitterionic PMPCs and cationic PDMAEMAs.

| **<u>Final Polymer</u>** | **<u>MPC</u>** |  | **<u>DMAEMA</u>** |  | **<u>ACVA</u>** |  | **<u>CPA</u>** |  |
| --- | --- | --- | --- | --- | --- | --- | --- | --- |
|  | *mass (g)* | *mols* | *mass (g)* | *mols* | *mass (mg)* | *mols* | *mass (mg)* | *mols* |
| *PMPC<sub>15</sub>* | 1.118 | 0.004 |  |  | 23.579 | 8.415E-05 | 70.530 | 2.525E-04 |
| *PMPC<sub>25</sub>* | 1.118 | 0.004 |  |  | 14.147 | 5.049E-05 | 42.318 | 1.515E-04 |
| *PMPC<sub>35</sub>* | 1.118 | 0.004 |  |  | 10.105 | 3.606E-05 | 30.227 | 1.082E-04 |
| *PDMAEMA<sub>80</sub>* |  |  | 4.950 | 3.149E-02 | 7.352 | 2.624E-05 | 21.992 | 7.872E-05 |
| *PDMAEMA<sub>160</sub>* |  |  | 3.960 | 2.519E-02 | 2.941 | 1.050E-05 | 8.797 | 3.149E-05 |

### 4.4. Synthesis of the Amphiphilic Block-Copolymers

The block-copolymers (PDMAEMA-b-PMPC) were produced by chain-extending the PMPC chain transfer agent via RAFT polymerization with DMAEMA monomers. Different degrees of polymerization (DP_x_) were targeted, with different cationic segment lengths (PDMAEMA). As a result, two different block-copolymers were synthesized, altering the ratio between PDMAEMA and PMPC ([PDMAEMA]/[PMPC] _x_) (where x is equal to 5 and 10). ACVA was used as an initiator in a molar ratio of 1/3 to PMPC, like PMPC and PDMAEMA synthesis. The reactions were performed in methanol (10%, w/v) at 60 °C for 24 h. Finally, the block-copolymers were purified by dialysis with 12-15 KDa membrane at 4°C overnight. Table 2 contains the reagent amounts.

**Table 2.** Reagents used to synthesize block-copolymers PDMAEMA-b-PMPC.

| **<u>Final Polymer</u>** | **<u>PMPC</u>** |  | **<u>DMAEMA</u>** |  | **<u>ACVA</u>** |  |
| --- | --- | --- | --- | --- | --- | --- |
|  | *mass (g)* | *mols* | *mass (g)* | *mols* | *mass (g)* | *mols* |
| *PDMAEMA-b-PMPC<sub>5:1</sub>* | 0.100 | 5.95E-06 | 4.67E-03 | 2.974E-05 | 5.56E-04 | 1.983E-06 |
| *PDMAEMA-b-PMPC<sub>10:1</sub>* | 0.100 | 5.95E-06 | 9.35E-03 | 5.947E-05 | 5.56E-04 | 1.983E-06 |

### 4.5. IONP Coating by Block-Copolymer PDMAEMA-b-PMPC

The IONPs were coated by block-copolymer (PDMAEMA-b-PMPC), mixing both solutions with a syringe needle without requiring robust equipment. The final solution was aspired and released by a 1 mL syringe equipped with a needle (0.55 x 20 mm) 10 times. This process with vigorous turbulence was performed to promote block-copolymer deposition over the IONP surface, forming the core-shell structure. Two different proportions (w/w) were tested, being IONP: block-copolymer (IONPP) equals 1:5 and 1:10, and IONP and block-copolymers were diluted in water.

### 4.6. Aptamer functionalization

To achieve the final nanoparticle structure called IONPPPs, IONPs coated by block-copolymer (IONPP) were functionalized with aptamer. Functionalization was performed by adding HER2 aptamer (2µM) conjugated in the 5’ with FAM fluorescent molecule on IONPP. First, the aptamers were heated at 90°C for 5 min and then cooled to room temperature over 15 min. The proportion was 1:60 (HER2 aptamer: IONPP, w/w). The reaction was performed in room temperature for 30 min under magnetic stirring.

### 4.7. Average diameter (z-average) / ζ-potential

All nanoparticle structure size and ζ-potential distributions were measured via dynamic light scattering using a Zetasizer Nano ZS90 μV (Malvern) at 25°C. The samples were diluted in ultrapure water. The measurements were performed in triplicate and analyzed using Zetasizer software, with an automatic titrator for pH assays.

### 4.8. X-ray diffractometry (XRD)

XRD was performed in a Bruker D8 Advance diffractometer to identify the crystal phases present in the synthesized samples. The IONP suspension was dried, powdered in a mortar, and compacted in a glass sample holder. The X-ray patterns were collected between 20 and 80° in the 2θ range using Cu Kα radiation, with 1.000 W (40 kV, 25 mA).

### 4.9. Thermogravimetric analysis (TGA)

TGA was performed in TGA/DSC (Mettler Toledo) equipment, in a 10 °C min^−1^ heating rate from 25 to 1000 °C, nitrogen atmosphere at 50 mL min^−1^ flow rate.

### 4.10. ^1^H Nuclear magnetic resonance spectroscopy (NMR)

To record ^1^H NMR spectra of each sample, approximately 10-20 mg were dissolved in 0.6 mL of CD_3_OD. The NMR spectra were recorded on the Agilent DD2 spectrometer from the Brazilian Biosciences National Laboratory (Brazilian Center for Research in Energy and Materials CNPEM), operating in Larmor frequency of 499,726 equipped with a triple resonance probe. Chemical shifts for protons were reported in parts per million (ppm) downfield from tetramethylsilane (TMS, 0 δ) and were referenced to the residual proton in the NMR solvent (CD_3_OD: 3.30 ppm).

### 4.11. Vibrating Sample Magnetometer (VSM)

The magnetic properties of the IONP and IONPP were evaluated using a conventional vibrating sample magnetometer (VSM 4500 EG&G). The samples were dried and introduced into an empty drug capsule before being inserted into the equipment. The measurements were performed at room temperature under an applied magnetic field ranging from −20 kOe to +20 kOe.

### 4.12. Fourier transform infrared spectroscopy (FT-IR)

For FT-IR analysis, Nicolet iS5/iD3 (ATR) equipment was used with wavenumbers from 600 to 4000 cm-1. The measurements were performed using dried samples.

### 4.13. Transmission Electron Microscopy (TEM)

The particle shape and size distribution of IONP and IONPP were examined by TEM using a JEOL JEM210F0 microscope operating at 200 kV and equipped with CM200 FEG (Philips) camera. The samples were prepared by water suspension dilution and dropped onto copper grids coated within holey carbon ultra-thin car film, 400 mesh (#01824, Ted Pella, EUA). The grids were treated in EasiGlow (I) (Ted Pella, EUA) at 25 mA for the 50s. The size distribution, average particle size (DTEM), and standard deviation (SD) were determined by measuring at least 150 nanoparticles using the free software ImageJ version v1.53t.

### 4.14. Nanoparticle Tracking Analysis (NTA)

NTA measurements of the EVs were performed using a NanoSight NS300 (NanoSight, Amesbury, United Kingdom) equipped with a sample chamber with a 532 nm green laser. The samples were measured at room temperature in PBS for 60 s with camera gain adjustments appropriate for the correct focusing of the EVs. The software NTA 3.4Build 0013 was used for data capture and analysis. Nanoparticle sizing and concentrations were determined based on the Brownian motion of individual particles and light scattering measurements.

### 4.15. Gel retardation assay

Gel electrophoresis was performed to evaluate the functionalization of (co)polymers and aptamers. The polymer-aptamer nanocomposites were prepared at different w/w ratios from 0.5 to 60.0 and loaded onto 3% agarose gels with glycerol 50% (2 μL). The analysis occurred in 1 × Tris-acetate-EDTA (TAE) buffer at 100 V for 40 min in a 15 cm electrophoresis chamber length. The bands were visualized using Carestream Gel Logic GL2200 Imaging System equipment.

### 4.16. Cell culture

MDA-MB-231 (ATCC HTB-26), SKBR3 (ATCC HTB-30), and HaCaT cells (ATCC CRL-4048) were maintained in RPMI 1640 medium supplemented with 10% fetal bovine serum (FBS) at 37°C under 5% CO_2_ in a humidified atmosphere.

### 4.17. Cytotoxicity assays

For the MTT assay, MDA-MB-231 cells were grown in 96-well plates at an initial density of 10000 cells/well in 100 μl medium and incubated for 24 h before adding (co)polymers. The cytotoxicity polymers were tested with different concentrations from 0.5 to 150.0 µg/mL. Each concentration was analyzed at least in triplicate. After additional incubation for 48 h, the medium was replaced with 200 μl serum-free medium, and 20 μL MTT solution (5 mg/mL) was added per well. After 4 h, 100 μL of DMSO was added. The plate was agitated for 10 min, and the absorbance of each well was performed on a PerkinElmer EnSpire 2300 multilabel plate reader at 492 nm.

For the fluorescent cytotoxicity assays, HaCaT cells were seeded at a density of 5000 cells/well in 384-well plates in RPMI medium supplemented with 10% of fetal bovine serum. After seeding, the cells were incubated with serial dilutions of PMPC_35_, PDMAEMA_80_, and PDMAEMA-b-PMPC_10:1_. The cells were incubated with 1 µg/mL propidium iodide (PI) (Sigma-Aldrich), MitoTracker probe (Life Technologies) 500 nM and 5 µg/mL Hoechst 33342 for 40 min in serum-free medium, at 37 °C, 5% CO_2_. The positive death control used was arsenite 2mM. Cells were evaluated using fluorescence microscope and plate reader Operetta (PerkinElmer) and analyzed with Columbus v2.8 software. GraphPad Prism (GraphPad Software, USA) version 8.0.1 was used for all statistical tests. The viability was normalized by the untreated cells, while nucleus volume and mitochondrial mass were evaluated by adjustment of inhibitor dose response curves using the log(inhibitor) vs. response function by the program GraphPad Prism (GraphPad Software, USA).

### 4.18. Fluorescence microscopy

MDA-MB-231 and SKBR3 cells were seeded at 5000 cells/well density in 96-well plates in RPMI medium supplemented with 10% fetal bovine serum. After 24 h, the block copolymer-aptamer complex (HER2 aptamer 1µM) was added with a final incubation time of 48 h. Then, the medium was replaced with PBS, and cells were evaluated using fluorescence microscope and plate reader Operetta (PerkinElmer) and further analyzed with Columbus v2.8 software.

### 4.19. Flow Cytometry

The MDA-MB-231 and SKBR3 tumor cells (2 x 10^5^) were incubated with IONPPP (1 µM HER2 aptamer) for 30 min at 37°C in the binding buffer (Hanks Buffer) (Z. Liu et al., 2012). Then, the samples were washed (400 rpms for 5 min at 4°C) once with 200 µl of binding buffer and resuspended in 200 µl PBS. For the magnet^+^ assay, after incubation, we added the magnet to the lateral microtube for 30 min at 37°C. After that, the supernatant was removed, and the complex was washed with the binding buffer (400 rpms for 5 minutes at 4°C). The sample acquisition was performed using BD FACS Canto II flow cytometer, and the FlowJo software (version 10) was used for analysis.

## 5. Results

### 5.1. Iron Oxide Nanoparticle Characterization

The co-precipitation method was applied to synthesize the iron oxide nanoparticles, which demand Fe^3+^ and Fe^2+^ mixed in a basic solution, though a green and easier-to-scale-up method. The XRD (Fig. 1A) results demonstrate peaks at 2θ = 30.2°, 35.5°, 43.2°, 53.5°, 57.1°, and 62.9° corresponding, respectively, to (220), (311), (400), (422), (511), and (440) crystalline planes of magnetite phase (JCPDS 190629). These results agree with ATR-FT-IR spectra (Fig. 1B). The FT-IR spectrum shows absorption bands at 3400, 2400, 1600, 1450, 1200, 630, and 580 cm^-1^. These observed bands are probably due to O-H (3400, 2400, 1600 cm^-1^) stretching, Fe-O (630 and 580 cm^-1^) stretching, and S=O (1200 and 1450 cm^-1^) stretching related to sulfate ions adsorbed in the surface of the nanoparticles.

**Figure 1.**
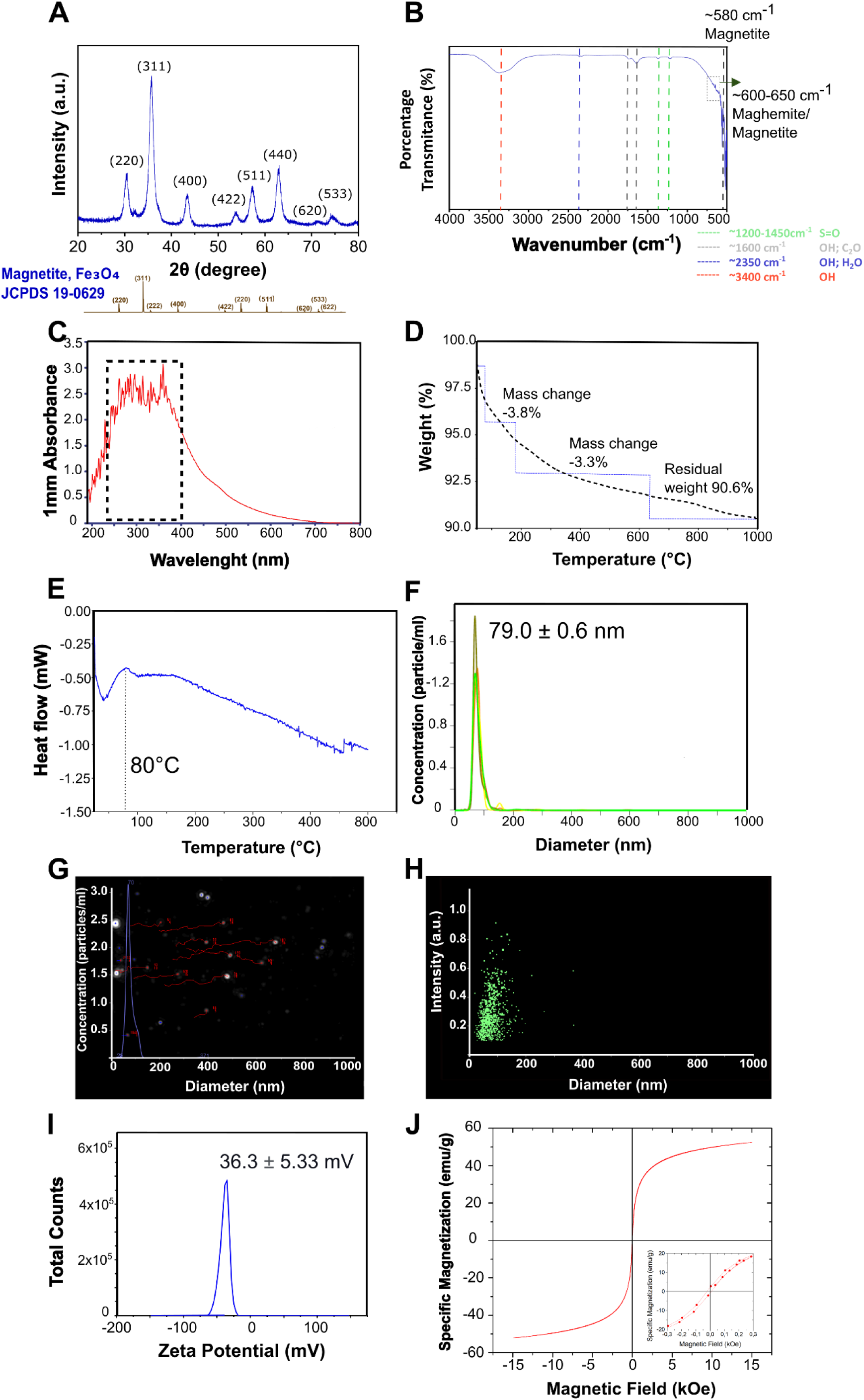
IONP Characterization. **A)** XRD (X-ray diffraction), **B)** FT-IR (Fourier-transformation infrared spectroscopy), and **C)** UV-VIS (Ultraviolet-visible spectroscopy) results related to the chemical composition characterization of IONPs; **D)** TGA (Thermogravimetric Analysis) results about the thermal decomposition and **E)** DSC (Differential Scanning Calorimetry) thermal events results; **F-H)** NTA (Nanoparticle Tracking Analysis), z-average size in water solution; **I)** ζ-Potential results in water solution by DLS (Dynamic light scattering), and **J)** VSM (Vibrating Sample Magnetometer) magnetic IONP results.

We also evaluated the optical properties of IONP in the UV-VIS spectrum (Fig. 1C). The results demonstrate a maximum absorption area between 250 and 300 nm. Finally, thermogravimetry analysis (TGA) was realized until 1000°C in a nitrogen atmosphere (Fig. 1D). We observed a loss of water molecules first (3.8% at 20-120°C), with 3.3% mass loss until 346°C and 2.3% mass loss until 1000°C, resulting in a total mass loss of around 9.4%. In addition, the DSC results demonstrated one endothermic event at 80°C with thermal decomposition until 500°C (Fig. 1E).

The hydrodynamic size of IONP in water solution was 78.04 ± - 0.4 nm by NTA (Fig. 1F-H). Figure 1G demonstrates a typical NTA video frame, while Fig. 1H shows the size distribution by scattergrams, revealing the highest particle concentration at lower intensity values. The ζ-potential of -36.3 ± 5.33 mV (Fig. 1I). The superparamagnetic property was confirmed by the highly reversible hysteresis curves measured in the VSM (Fig. 1J), with a saturation magnetization value (M_s_) of 60 emu/g. The weak coercivity showed by the sample (∼30 Oe) can be attributed to a small fraction of magnetically blocked nanoparticles.

The morphology and size of IONPs were verified by TEM (Fig. 2). We observed a spherical nanoparticle (Fig. 2A-C) with an average 8.9 ± 1.9 nm diameter (Fig. 2D) and a polydispersity index (*σ* = SD/*D*TEM) of 0.2, suggesting a narrow size distribution in a monodisperse-like system (Fig. 2D). The SAED analysis also corroborates with the magnetite phase results (Fig. 2E) (Basavaraja et al., 2011; Farmer, 1976; Mishra et al., 2014). Finally, different planes can be observed in the HRTEM images of IONPs (Fig. 2F), with the following distances 2.1 Å (400), 2.4 Å (111), 2.9 Å (220), and 4.7 Å (111).

**Figure 2.**
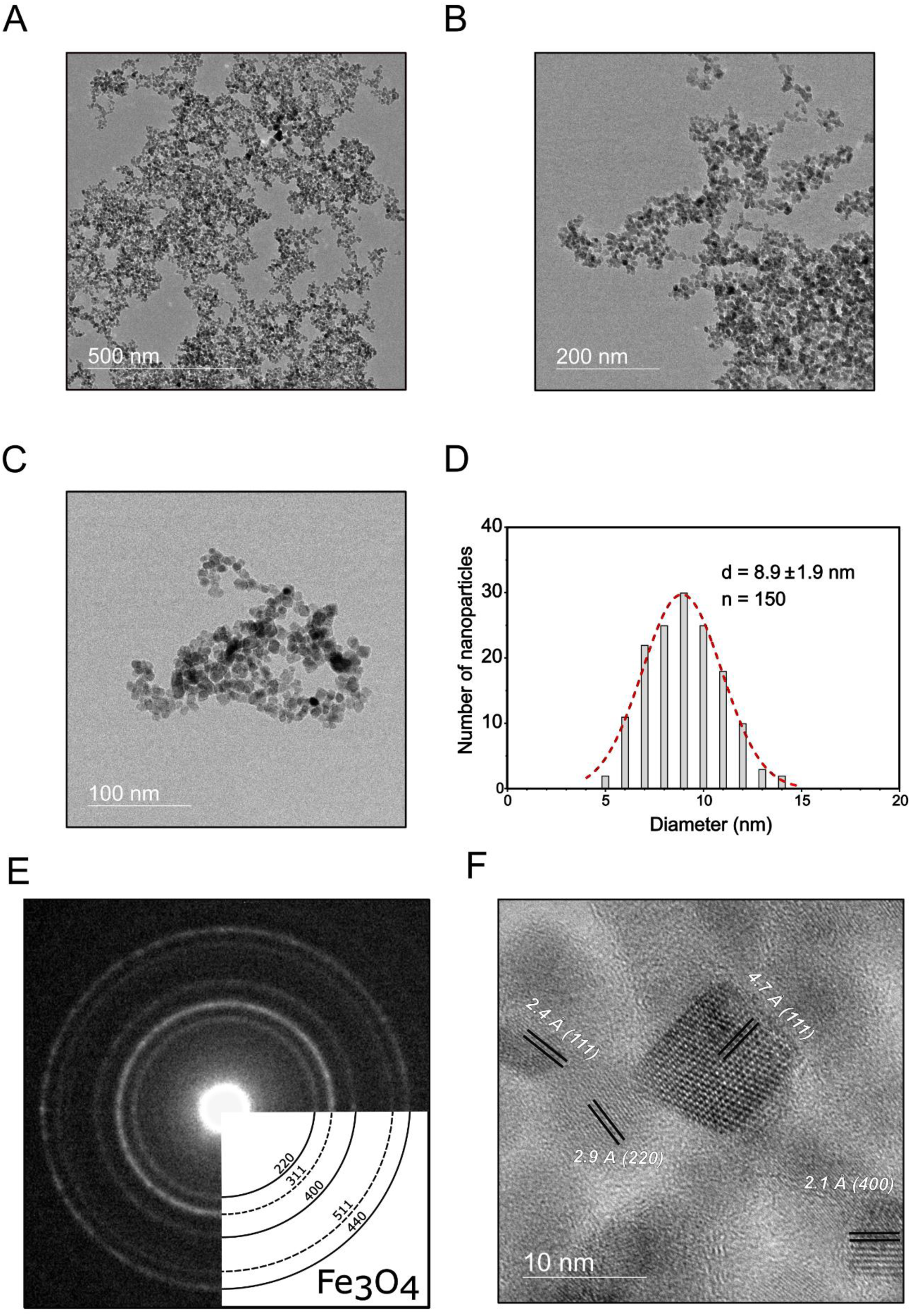
IONP Microscopic Characterization. MET/HRMET (High-resolution Transmission Electron Microscopy) images **A-C)** at three magnifications, **D)** frequency size distribution graph; **E)** crystallographic results by the SAED (Selected Area Electron Diffraction), and **F)** HRTEM image of IONPs with four lattice planes highlights.

The block-copolymers were used to coat the IONPs. The RAFT method synthesized all polymers and block-copolymers (Fig. S1A, S1D, and Fig. 3A). Firstly, we synthesized the macromonomers PMPC and PDMAEMA separately to understand the properties of each polymer. Then, we focused on two different proportions between the cationic and zwitterionic components of the block-copolymer PDMAEMA-b-PMPC. This step contemplated the theoretical molar proportions 5:1 and 10:1 between PDMAEMA and PMPC, forming, respectively, PDMAEMA-b-PMPC_5:1_ and PDMAEMA-b-PMPC_10:1_. Finally, ^1^H NMR analysis was performed to confirm the degree of polymerization (DP), the number-average molecular weight (M_n_), and the residual monomer of each synthesis. All results related to macromonomers PMPC and PDMAEMA to ^1^H NMR spectra are described in Table 3, and details of each synthesis are in the method section. Our results demonstrated no residual monomer to PDMAEMA synthesis (Fig. S2A) and PMPC after purification (Fig. S2B). PMPC demonstrated the experimental DP above the theoretical DP, while PDMAEMA showed beyond the experimental DP (Table 3). Moreover, Fig. S2C represents the essential assigned signals from the cationic (L’ and N’, for PDMAEMA) and the zwitterionic (E’, F’, D’, G’, for PMPC) components in the ^1^H NMR results of the block-copolymers for all three proportion syntheses.

**Figure 3.**
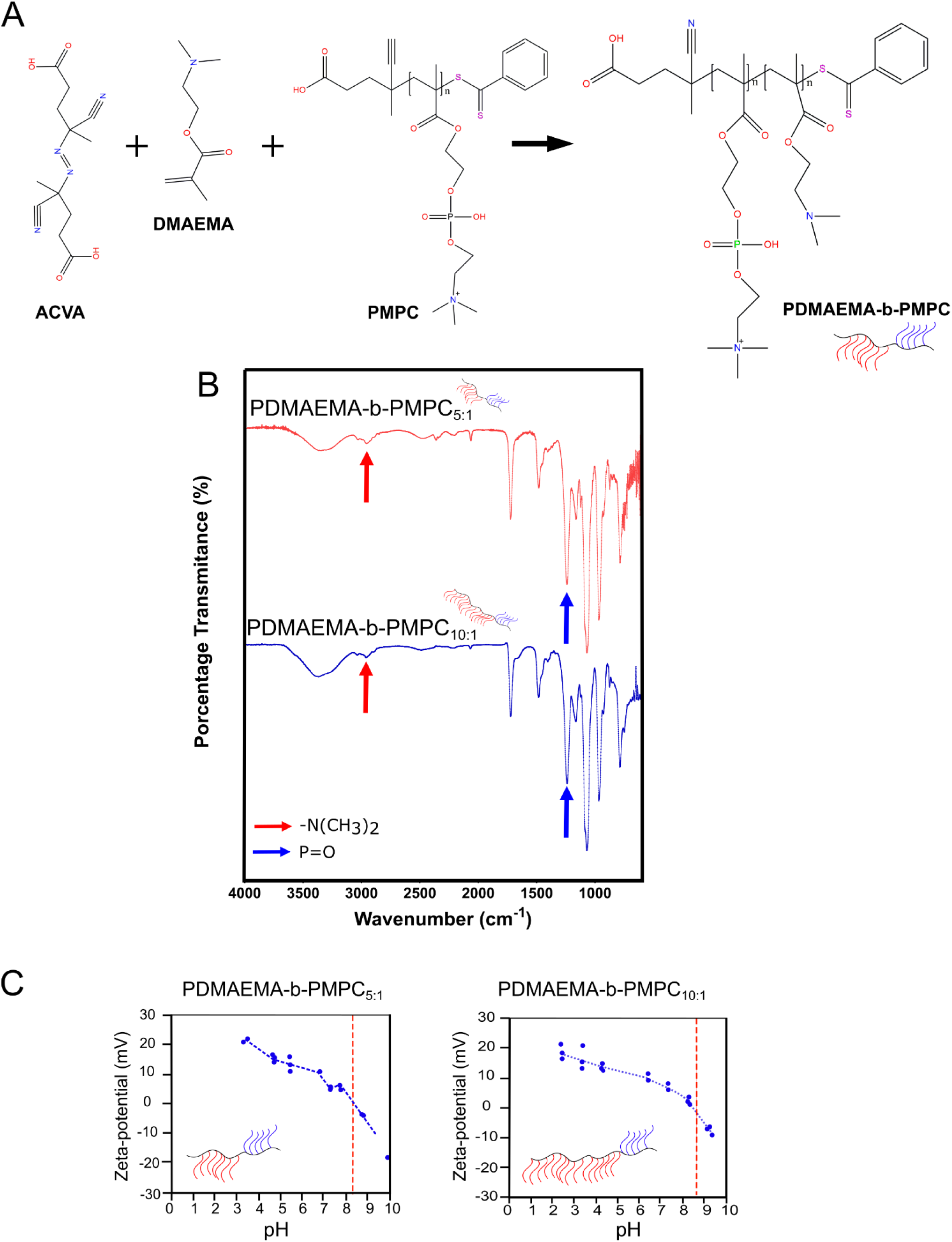
Block-Copolymers characterization. **A)** Scheme representation of experimental approach synthesis of the block-copolymer PDMAEMA-b-PMPC by RAFT method; **B)**; FT-IR (Fourier transformation infrared spectroscopy) spectrum of the PDMAEMA-b-PMPC_5:1_ and PDMAEMA-b PMPC_10:1_; **C)**; z-average size and ζ-Potential in broad pHs of the PDMAEMA-b-PMPC_5:1_ and PDMAEMA b-PMPC_10:1_ in water solution by DLS (Dynamic light scattering).

**Table 3.**
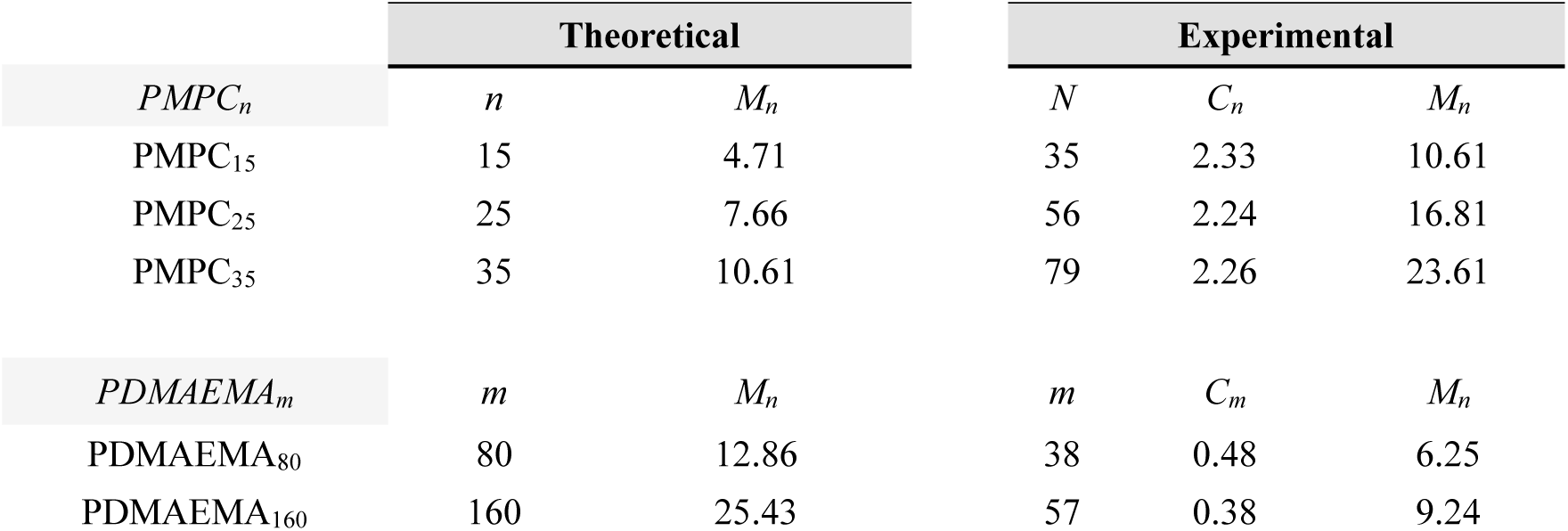
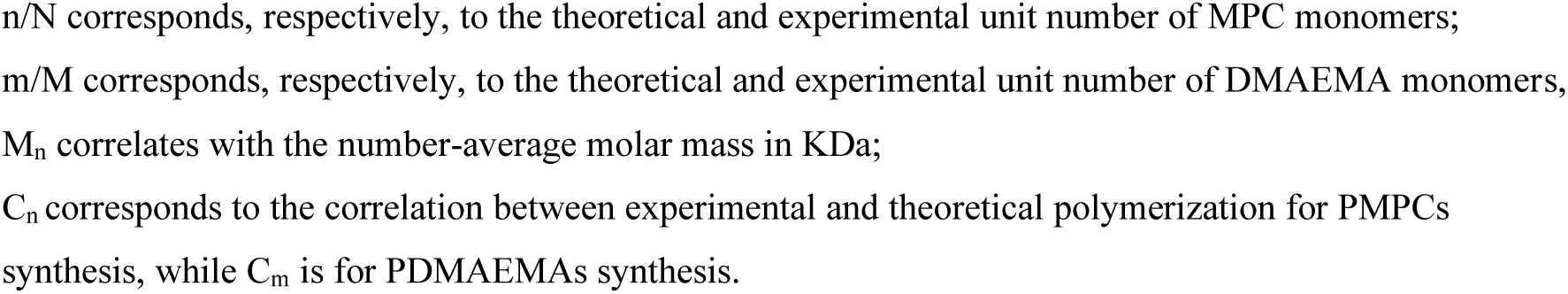
Degree polymerization (DP) of macromonomers PMPC and PDMAEMA.

|  | Theoretical |  | Experimental |  |  |
| --- | --- | --- | --- | --- | --- |
|  | *n* | *M<sub>n</sub>* | *N* | *C<sub>n</sub>* | *M<sub>n</sub>* |
| *PMPC<sub>n</sub>* |  |  |  |  |  |
| PMPC<sub>15</sub> | 15 | 4.71 | 35 | 2.33 | 10.61 |
| PMPC<sub>25</sub> | 25 | 7.66 | 56 | 2.24 | 16.81 |
| PMPC<sub>35</sub> | 35 | 10.61 | 79 | 2.26 | 23.61 |
| *PDMAEMA<sub>m</sub>* | *m* | *M<sub>n</sub>* | *m* | *C<sub>m</sub>* | *M<sub>n</sub>* |
| PDMAEMA<sub>80</sub> | 80 | 12.86 | 38 | 0.48 | 6.25 |
| PDMAEMA<sub>160</sub> | 160 | 25.43 | 57 | 0.38 | 9.24 |

*n/N* corresponds, respectively, to the theoretical and experimental unit number of MPC monomers; *m/M* corresponds, respectively, to the theoretical and experimental unit number of DMAEMA monomers, *M<sub>n</sub>* correlates with the number-average molar mass in KDa; *C<sub>n</sub>* corresponds to the correlation between experimental and theoretical polymerization for PMPCs synthesis, while *C<sub>m</sub>* is for PDMAEMAs synthesis.

ATR-FT-IR was also realized to confirm the composition of polymers and the PDMAEMA-b-PMPC. The PDMAEMA shows bands at ∼3000 cm^-1^ referents to C-H stretching in ν(N(CH_3_)_2_), 1700 cm^-1^ (C=O), and the 1400-1500 cm^-1^ bands are related to methylene groups (Fig. S1B). The PMPC spectrum (Fig. S1E) presents bands at 1240 and 1056 cm^-1^ related respectively to symmetric and asymmetric stretching of the ν_as,s_ (P-O) and ν_as,s_ (P−O), while the quaternary amine ν(N+(CH_3_)_4_) is represented at 958 cm^-1^ and the carboxylate group (ν(COO−)) at 1727 cm^-1^. The block-copolymer structuration shows the P-O and -N(CH_3_)_2_ stretching, 1240 cm^-1^, and 3000 cm^-1^ bands, respectively (Fig. 3B). The hydroxyl (O-H) stretching is observed at 3400 cm^-1^ in all (co)polymers synthesized.

We also measured the ζ-potential of nanoparticles to confirm the block-copolymer synthesis, whereby more PDMAEMA component means more cationic total charge in the block-copolymer due to the protonation of (N(CH_3_)_2_) groups. Fig. 3C. clearly illustrates the shift to achieving ζ-potential of block-copolymers equals zero with a higher PDMAEMA component. Furthermore, the ζ-potential of PDMAEMA macromonomers demonstrated positive ζ-potential in acid and neutral pH (Fig. S1C). In contrast, PMPC remains negative in all pH analyses (Fig. S1F), with a more negative charge related to the polymer in basic pH.

We then evaluated the metabolic viability of cells after incubation with polymers and block-copolymers by MTT assay (Fig. S3A-B), as well as the relative number of live cells (Fig. S3C), necrotic cells (as measured by propidium iodide PI – incorporation) (Fig. S3D), mitochondrial mass (Fig. S3E), and nucleus volume (Fig. S3F). This way, we evaluate the toxicity of polymers and block-copolymers synthesized by different biological approaches a) MTT assay measures the cells metabolic activity, indicating the metabolic viability of cells b) PI incorporation verifies the membrane integrity, resulting in the number of necrotic cells; c) differences in mitochondrial mass indicate an alteration in energy production by mitochondria, and d) alteration in nucleus volume demonstrates cellular damage, indicating genotoxicity. No relevant reduction in cell viability was observed for PDMAEMA and PMPC in the MTT assay (Fig. S3A). However, the block copolymer showed only a minimal decrease in cell viability, as assessed by the MTT assay, with a maximum reduction of 20% (Fig. S3B). Furthermore, the toxicity related to cell number (Fig. S3C), cell death (Fig. S3D), mitochondrial mass (Fig. S3E), and nucleus volume (Fig. S3F) did not significantly change compared to the death control, regardless of the tested polymer molecule.

### 5.3. IONP Coating by Block-Copolymer (IONPP)

We studied two different proportions (weight/weight) between IONP and block copolymers: 1:5 and 1:10. In the proportion 1:5. We detected precipitation in 1 h (data not showed); in this regard, we followed only the IONPP_1:10_ ratio in the next steps. To confirm the encapsulation of IONP by block block-copolymer PDMAEMA-b-PMPC, we first verified the z-average size (Fig. 4B) and ζ-potential of the final nanostructure (Fig. 4C). While uncoated IONP demonstrated a size of 208.1 ± 1.5 nm, IONPP presented a size of 281.7 ± 1.53 nm (Fig. 4B); both groups showed a low polydispersity index (< 0.2) (Fig 4B). Furthermore, while the block-copolymer alone had a positive value (21.57 ± 1.429 mV), we measured a change in the total surface charge from -34.50 mV (IONP) to 3.99 mV (IONPP) (Fig. 4C). Importantly, IONPP_1:10_ maintained the superparamagnetic behavior (Fig 4E).

**Figure 4.**
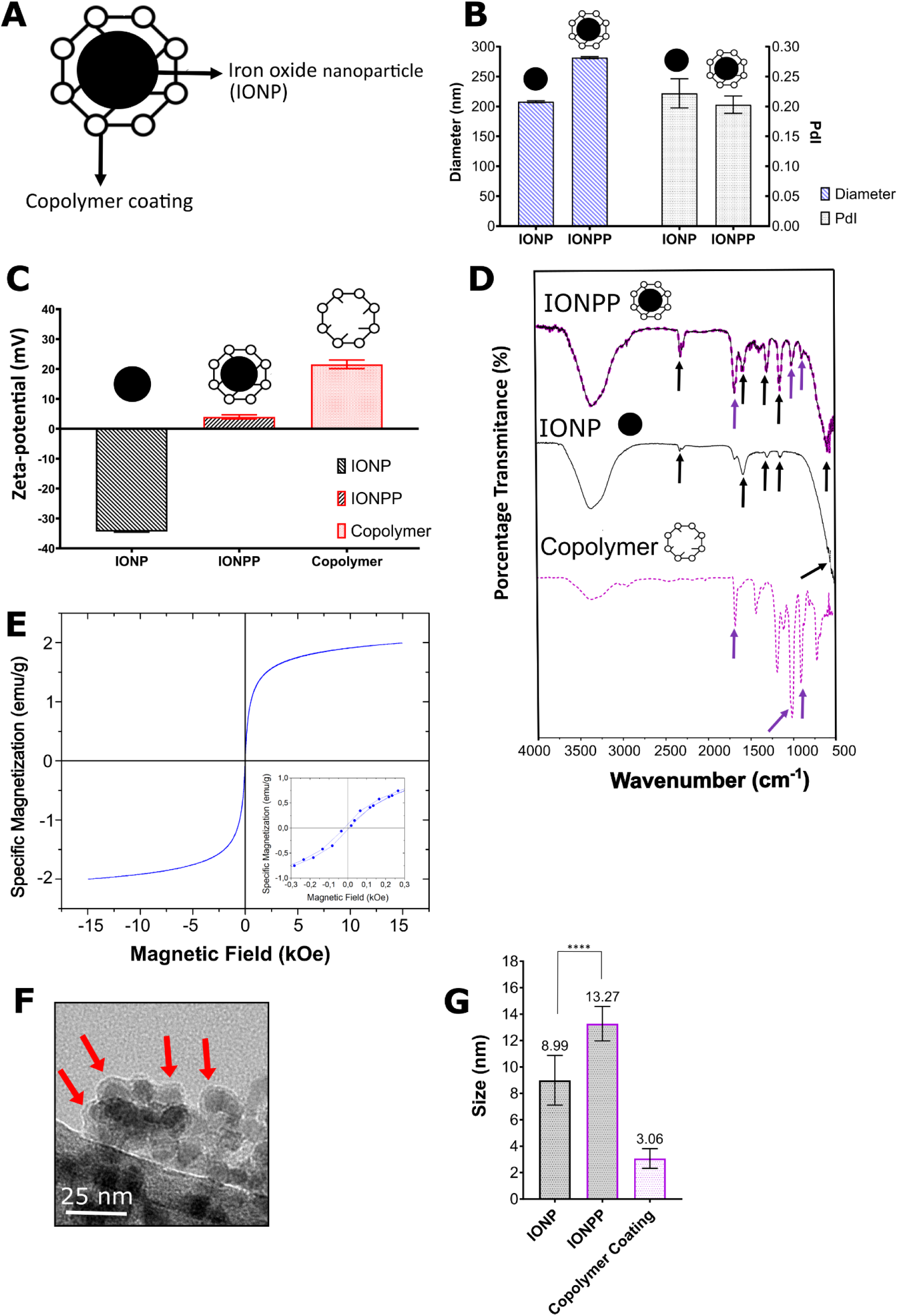
IONP coating by the block-copolymer (IONPP). **A)** Scheme representation of IONPs coated by the block copolymer PDMAEMA-b-PMPC; **B)** z-average size data and PdI of IONP and IONPP by DLS (Dynamic light scattering); **C)**; ζ-Potential data of IONP, IONPP, and block-copolymer by DLS; **D)** FT-IR (Fourier-transformation infrared spectroscopy) spectrum of IONP, IONPP, and block-copolymer (PDMAEMA-b-PMPC), black arrows indicated IONP characteristics band, while purple arrows showed the block-copolymer characteristics bands; **E)**; IONPP VSM (Vibrating Sample Magnetometer) results; **F)** TEM (Transmission Electron Microscopy) image of IONPP, the red arrows indicating the IONPP structures, and **G)** size of IONP, IONPP, and block-copolymer coating thickness. **** p-value < 0.0001; unpaired t-test. Scale bar = 25 nm.

As expected, the IONPP morphology was spherical (Fig. 4F) with a distribution size of 13.27 ± 1.30 nm (Fig. 4G). It was possible to see the metal component in the middle of IONPP (red arrow) by TEM (Fig. 4F). Furthermore, the block-copolymer coating has 3.1 ± 0.75 nm (Fig. 4G).

Then we evaluated the binding between IONP and block-copolymers after using magnet to capture the IONPP (IONPP_1:10_ ^magnet+^) (Fig. 5A). The size of IONPP_1:10_ ^magnet+^ was 992.5 ± 40.43 nm in an aqueous solution, with a higher aggregation than IONPP_1:10_ ^magnet-^ (Fig 5B). However, after being sonicated for 1 h, the IONPP_1:10_^magnet+^ reduced to 337.4 ± 3.837 nm (Fig. 5B, the red arrow). The IONPPP_1:10_^magnet+^ ζ-potential was positive after using a magnet (14.9 ± 0.3 mV) (Fig. 5C). This result demonstrated that the block copolymer maintains binding to IONP. However, we observed higher values of ζ potential on the IONPP_1:10_ ^magnet+^, which can be explained by the aggregation of NPs, with more block-copolymer on the surface of IONP. We also evaluated using FT-IR and TGA the IONPP_1:10_ ^magnet+^. The FT-IR results demonstrated bands related to block-copolymer 958 and 1240 cm^-1^ and IONP 630 cm^-1^ (Fig. 5D), the same result of IONPP_1:10_^magnet-^ (Fig. 4D). Furthermore, the TGA for IONPP_1:10_ ^magnet+^ shows a reduction mass of 8.30% until 181°C (Fig. 5E). The second thermal event finished around 522°C, with a residue of 36.40% of the mass. The last mass loss resulted in a final mass of 22.81%.

**Figure 5.**
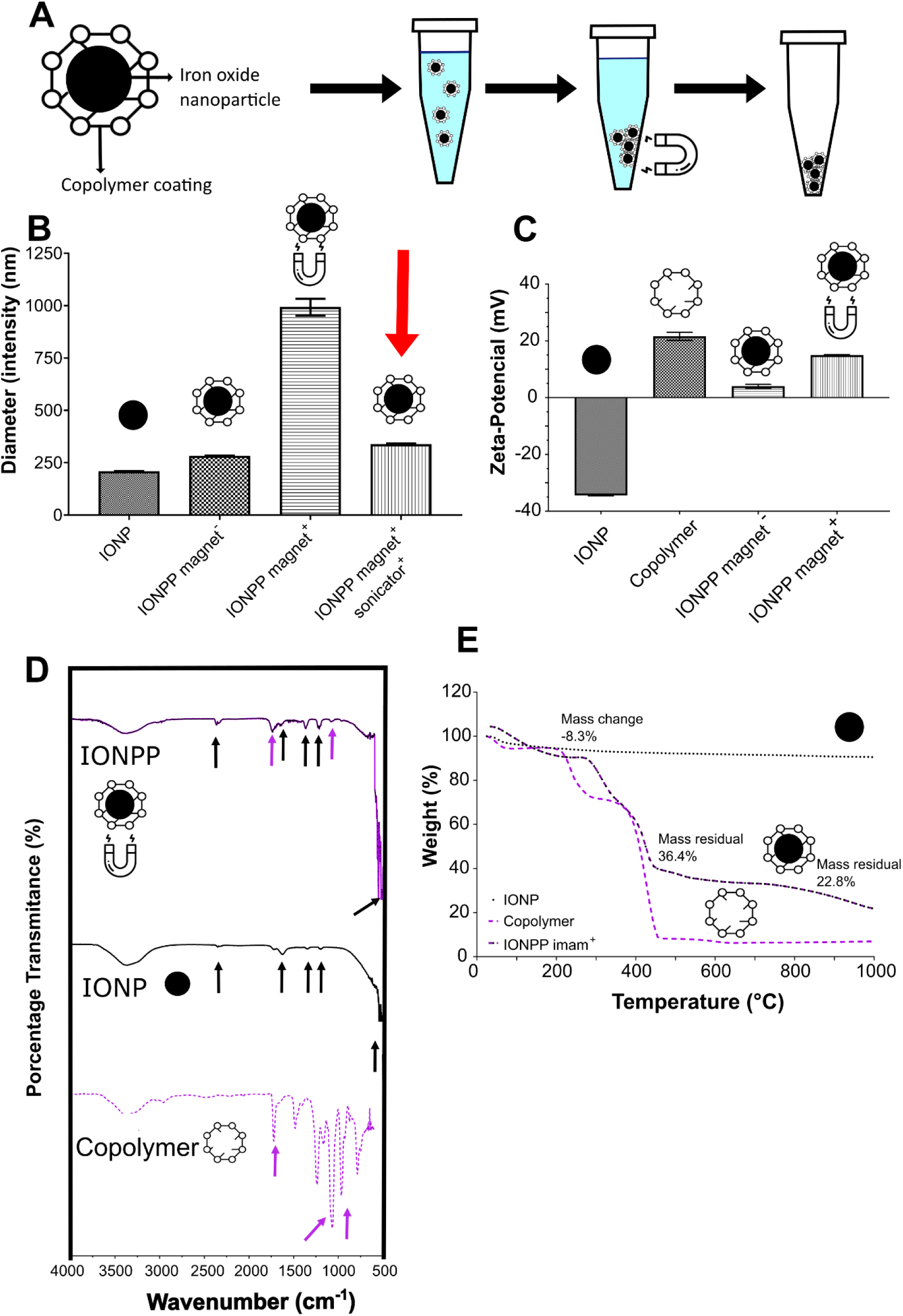
IONPP characterization after being captured by a magnet. **A)** Scheme representation of methodology employed for IONPP captured by magnet; **B)** z-average size of IONP and IONPP by DLS (Dynamic light scattering) before and after using magnet, the red arrow indicates the IONPP after using a magnet and the sonicator; **C)** ζ-Potential of the IONP, block-copolymer, IONPP before using a magnet, and results after IONPP captured; **D)** FT-IR (Fourier-transformation infrared spectroscopy) spectrum of IONP, IONPP, and block-copolymer (PDMAEMA-b-PMPC) after using a magnet, black arrows indicated IONP characteristics band, while purple arrows showed the block-copolymer characteristics bands; **E)** TGA (Thermogravimetric Analysis) results in weight (%/%) of IONP, block-copolymer, and IONPP after using the magnet.

Then, we evaluated the binding capacity of the block-copolymers to the anti-HER2 aptamer, as shown. By performing a gel retardation assay, we verified a more increased retardation of the aptamer the higher the amount of block-copolymer incubated (Fig. 6B). By measuring the migration distance of all gel-retarded bands. We calculated the EC_50_ of the block-copolymers to the aptamer as being 10.29 µg/mL (95% CI: 10.03 to 10.54 µg/mL) to PDMAEMA-b-PMPC_5:1_ and 7.07 µg/mL (95%CI: 6.89 to 7.25) to PDMAEMA-b-PMPC_10:1_ (Table 4). In comparison, the polymer PDMAEMA_80_ presented an EC_50_ of 0.79 µg/mL (95%CI: 0.34 to 1.38 µg/mL). The hydrodynamic size and the ζ potential of the block-copolymer PDMAEMA-b-PMPC_10:1_ aptamer complex was 237.2 ± 31.8 nm (Fig. 6D), showing a more compact size compared to the non-functionalized PDMAEMA-b-PMPC_10:1;_ the ζ-potential measured shifted from being positive (+18.7 ± 2.0 mV) without the aptamer to being negative (-2.3 ± 0.7 mV, Fig. 6E) in the presence of the nucleic acid.

**Figure 6.**
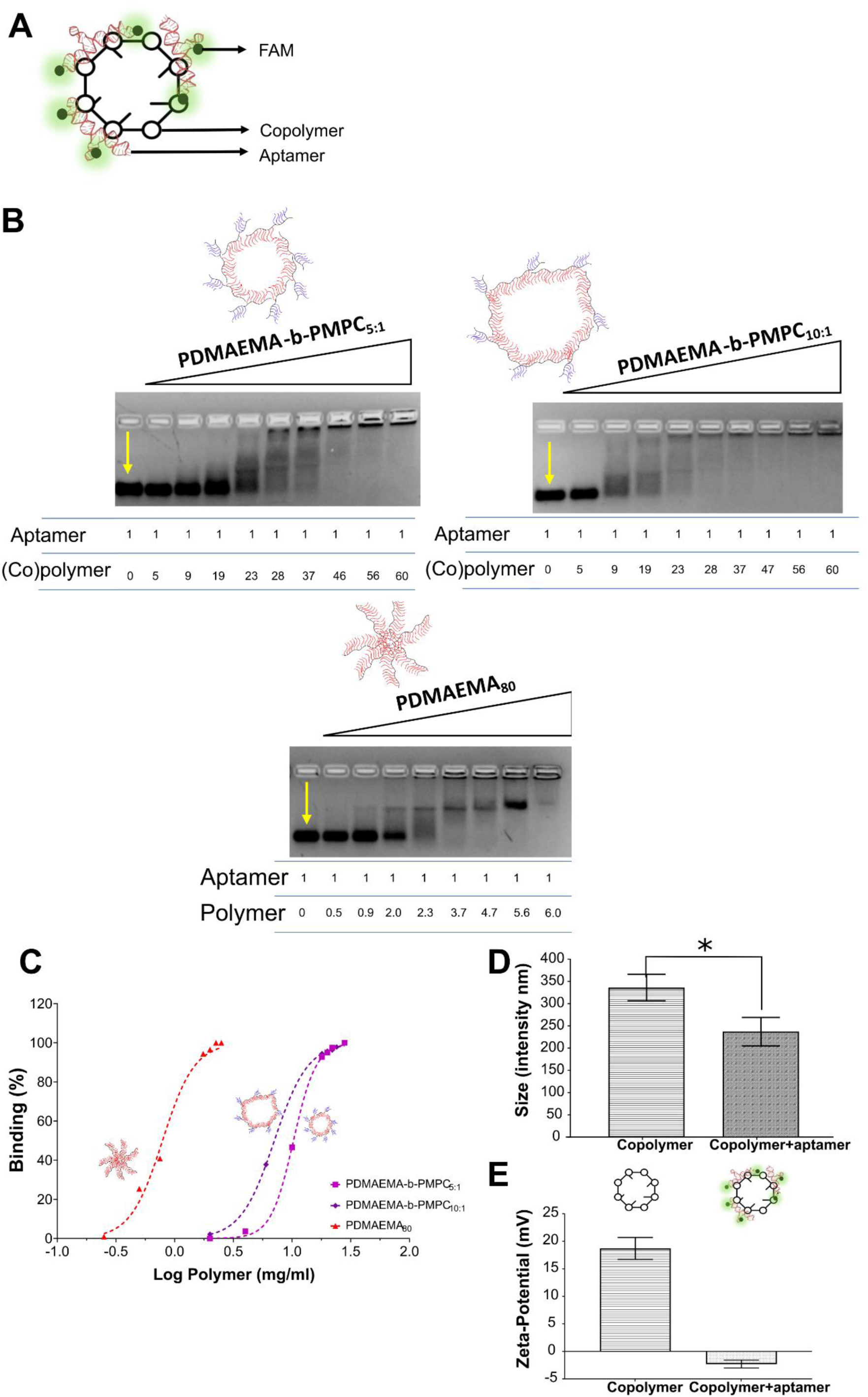
Functionalization binding results of block-copolymer PDMAEMA-b-PMPC and aptamers. **A)** Scheme representation of block-copolymer PDMAEMA-b-PMPC binding with aptamer, the block-copolymer-aptamer complex; **B)** Gel retardation assay; **C)** EC_50_ curves of the block-copolymer PDMAEMA-b-PMPC_5:1_, PDMAEMA-b-PMPC_10:1_, and the PDMAEMA_80_; the **D)** z-average size, and the **E)** ζ-potential results of block-copolymers (not) binding to aptamers. The yellow arrow shows the aptamer band without polymers; the numbers represent the proportion between polymer and aptamer (w/w). P-value *0.0171, unpaired t-test with Welch’s correction.

**Table 4.** The EC_50_, SD, and R² of the polymer-aptamer complex.

| <u>**Final Polymer**</u> | <u>**EC<sub>50</sub>**</u> | <u>**SD**</u> | <u>**R<sup>2</sup>**</u> |
| --- | --- | --- | --- |
|  | µg/mL | µg/mL |  |
| *PDMAEMA*<sub>80</sub> | 0.79 | 0.70 to 0.89 | 0.9912 |
| *PDMAEMA-b-PMPC*<sub>5:1</sub> | 10.29 | 10.02 to 10.55 | 0.9994 |
| *PDMAEMA-b-PMPC*<sub>10:1</sub> | 7.07 | 6.89 to 7.25 | 0.9997 |

We then evaluated the binding of IONPP (formed with PDMAEMA-b-PMPC_10:1_) to the anti-HER2 aptamer (IONPPP). The hydrodynamic IONPPP size was 297.2 ± 5.886 nm (Fig. 7B) with PdI 0.20 ± 0.04 (Fig. 7C), which demonstrated an increase compared to IONPP (287.4 ± 1.179 nm, Fig. 7B, Fig. S4A). The ζ-potential measured was 16.33 ± 0.25 mV, a smaller value than the IONPP (25.03 ± 0.38 mV, Fig. 7D, Fig S4B). To confirm that the aptamer was being complexed to the IONPP before and after magnet pull-down (Fig. 7E and 7F, respectively), we used SDS to break apart the molecules and analyzed them with agarose gel.

**Figure 7.**
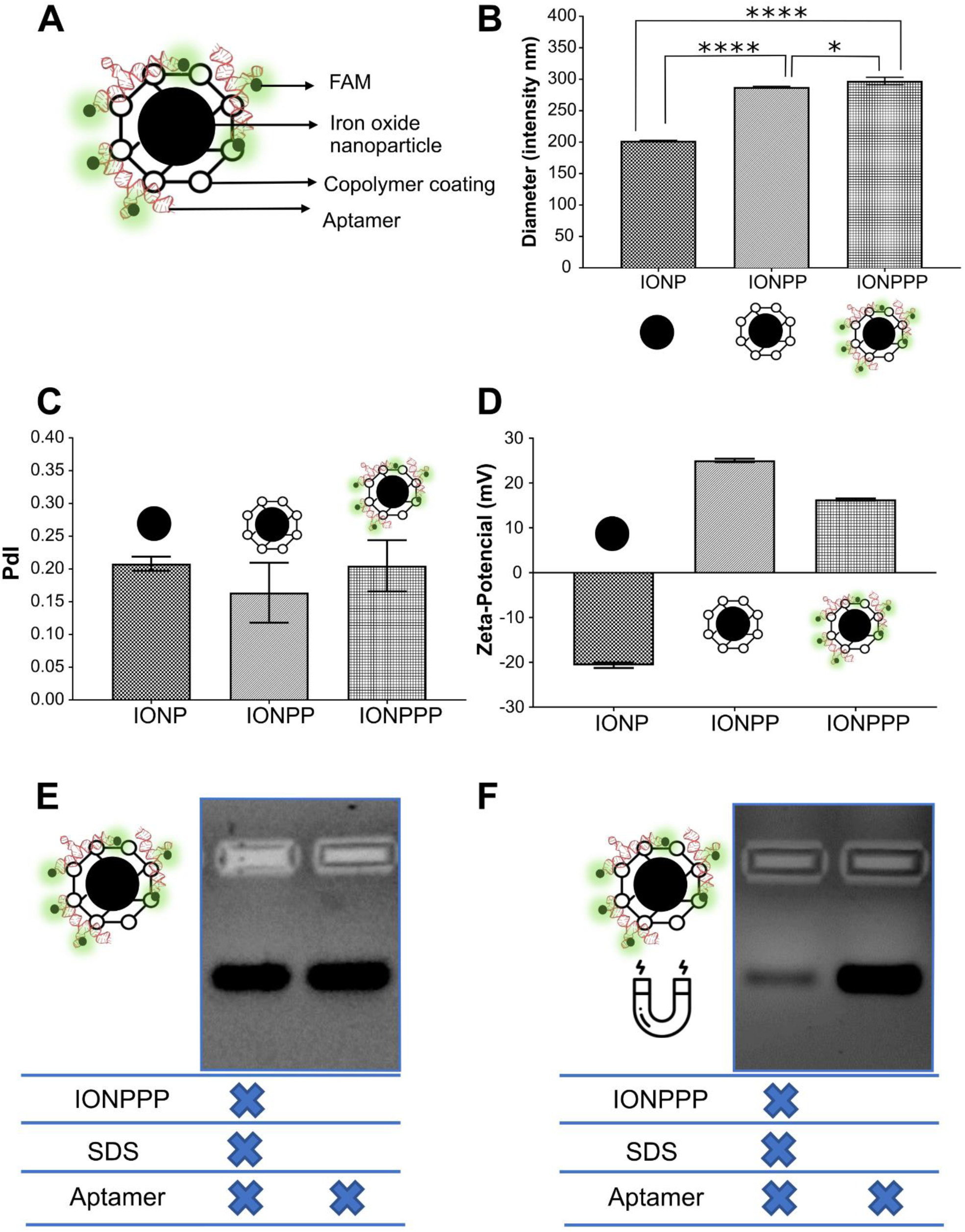
Iron oxide nanoparticles coated by block-copolymer and binding with aptamer (IONPPP) characterization. **A)** Scheme representation of iron oxide nanoparticles coated by the block copolymer PDMAEMA-b-PMPC and functionalized with aptamers; **B)** z-average size data of IONP, IONPP, and the IONPP by DLS (Dynamic light scattering) and **C)** the PdI results; **D)** ζ-Potential data of IONP, IONPP, IONPPP, and block-copolymer by DLS (Dynamic light scattering); **E)** gel retardation assay of IONPPP incubation with SDS without magnet, and **F)** after using magnet.**** p-value < 0.0001; * p value< 0.05 (0.0319); both unpaired t-test, Tukey’s multiple comparisons tests.

### 5.5. Isolation of HER2 breast cancer cells^+^ with IONPPP

The gate strategy in flow cytometry was employed to accurately identify and analyze SKBR3 (Fig. S5A) and MDA-MB-231 (Fig. S5B) cells bound to HER2-aptamer. Gates were set based on forward scatter (FSC) and side scatter (SSC) parameters to exclude debris and non-cellular particles. Subsequently, gating was performed to identify the FAM-5’HER2 Aptamer (FITC) for specific subset identification. Additionally, the pattern of IONP in a flow cytometer was verified (Fig. S5C) to exclude this area from further analysis.

We confirmed that the anti-HER2 aptamer exhibited increased recognition of the cell line SKBR3 (32.7%), which has a higher expression of HER2 (Mota et al., 2017), compared to the cell line MDA-MB-231 (7.2%) (Fig. 8A), which display lower levels of HER2 expression (Mota et al., 2017). This finding is consistent with our microscopy staining, which specifically occurred on the cell membrane SKBR3 (Fig S5D), as HER2 is a protein localized in the cell membrane (Mota et al., 2017). Next, we examined the autofluorescence of IONPP when incubated with both cells and observed less than 1% of positive cells (Fig. 8B). Subsequently, we assessed the binding capacity of the block copolymer (PDMAEMA-b-PMPC_10:1_) - aptamer complex to bind to the HER2^+^ SKBR3 cells, with 2.40-fold higher compared to the MDA-MB-231 cells (Fig. 8C).

**Figure 8.**
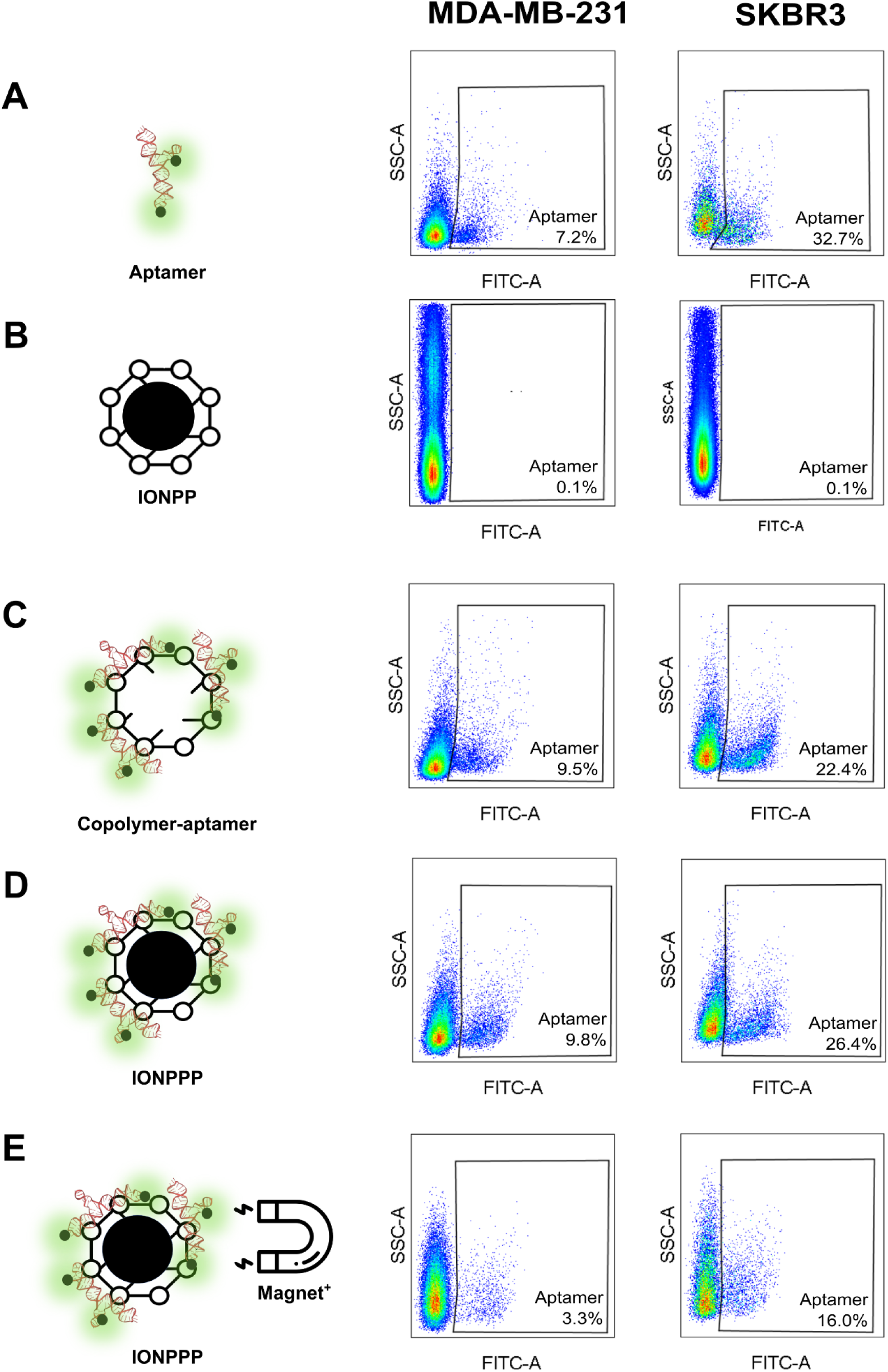
Application of IONPPP in tumor cells. Flow cytometry results in MDA-MB-231 and SKBR3 tumor cells of **A)** HER-2 Aptamer validation; **B)** IONPP autofluorescence results; **C)** the PDMMA-b PMPC – aptamer complex; **D)** IONPPP incubated with HER2 cell^-/+^ without using magnet IONPPP ^magnet^; and **E)** flow cytometry results of the IONPPP incubated with HER2 cell^-/+^ cell with magnet application (IONPPP ^magnet+^).

Furthermore, we proceeded to assess the binding capacity of IONPPP to SKBR3 cells compared to MDA-MB-231 cells. Following incubation with IONPPP, we observed that 9.8% of MDA-MB-231 cells exhibited fluorescence positivity. In contrast, the functionalized NP recognized 26.4% of SKBR3 cells (Fig. 8D). Next, we conducted a test to determine the number of positive cells that could be pulled down by a magnet after incubation with IONPPP magnet+. The results showed that 16.0% of SKBR3 cells were successfully captured by the magnetic NP. In contrast, only 3.3% of MDA-MB-231 cells were pulled down (Fig. 8E), which is 4.8-fold lower than the HER2^+^ SKBR3 cells. Despite the reduction in efficiency when using a magnet, our results demonstrated an increased sensitivity.

## 6. Statistical methods

All data are expressed as means ± standard deviations of a representative of at least three independent experiments carried out in triplicate. All analyses were performed using GraphPad Prism 8 Software, version 8.0.1. Statistical comparisons were made by the unpaired t-test, with Welch’s correction and Tukey’s multiple comparisons tests when applicable. In all statistical evaluations, p <0.05 was considered statistically significant.

## 7. Discussion

The careful planning of nanomaterials for biomedical purposes, especially in cancer treatment and diagnosis, necessitates thoroughly considering all constituent components functioning within a multifunctional framework. First, we synthesized an IONP using the classical co-precipitation method and confirmed the chemical composition using XRD, FT-IR, UV-VIS, TGA, and SAED techniques. XRD confirms the formation of iron oxide nanoparticles with a classical pattern of other studies (Cornell & Schwertmann, 2003b; Perecin et al., 2022) corresponding to an inverse spinel structure. However, XRD cannot detect the differences between maghemite and magnetite because these two phases are isostructural. The FT-IR spectra demonstrated the magnetite formation (580 cm^-1^ characteristic band) and some nanoparticle surface adsorbed molecules. These 1000-1200 cm^-1^ bands represent the sulfur group residue of the synthesis, which is observed in other studies (Perecin et al., 2021, 2022) and can be helpful to colloidal stabilization. Also, the bands between 600 and 650 cm^-1^ are associated with the Fe-O stretching related to the maghemite phase, indicating some oxidation of our IONPs. The SAED and HRTEM results corroborate with the chemical composition of the IONP, indicating a Fe_3_O_4_ cubic crystalline network, which is commonly found in the literature (Cornell & Schwertmann, 2003b).

IONPs absorb light in the UV region (Cornell & Schwertmann, 2003c). Therefore, they are strongly reflected in the visible/near IR regions, aligning with our results. The TGA results demonstrated the loss of water molecules and degradation of the adsorbed sulfate groups evidencing the thermal degradation pattern of IONPs. TEM and HRTEM verified the morphology, and we mainly observed a spherical shape. This morphology is expected due to low-temperature synthesis (less than 100°C), with probable Fe^2+^ excess over OH^-^, and it is corroborated by other studies (Cornell & Schwertmann, 2003d; Perecin et al., 2021). While the large size distribution is classical due to the synthesis method employed (Ajinkya et al., 2020; Ali et al., 2016; Beck et al., 2022). Our results demonstrated extensive and robust characterization of the IONPs, confirming the chemical composition aligned with the literature.

The IONPs have high aggregation characteristics when uncoated (Ali et al., 2016; Cornell & Schwertmann, 2003d, 2003a). Therefore, we used a biodegradable block copolymer PDMAEMA-b-PMPC to encapsulate the IONPs. PMPC was chosen to provide colloidal stability and better biocompatibility (Goda et al., 2015b; Perecin et al., 2022), while PDMAEMA binds to acid nuclei by electronic binding (C. Zhu et al., 2010). First, we synthesized and characterized the polymers separately and then the block copolymer. As a result, we observed a DP_experimental_ above of DP_theoretical_ for PMPC, also found in another study about PMPC synthesis by RAFT (Perecin et al., 2022). Furthermore, our results indicated a low monomer conversion of DMAEMA in PDMAEMA and the block-copolymer PDMAEMA-b-PMPC, with the same pattern as previous studies (L. Wang et al., 2019). The improvement to achieve precise DP for DMAEMA conversion could be tested in future studies by changing the proportion between initiator and macro-CTA, temperature, and polymerization time.

The FT-IR and ^1^H NMR results confirmed the PDMAEMA and the PMPC polymerization, such as the block-copolymer PDMAEMA-b-PMPC. For the zwitterionic and cationic polymers, we could observe the aromatic hydrogen signal in the CPA ring (carbons A’, B’, and C’), such as the hydrogen corresponding to the MPC molecule (D’, E’, F’, and G’) and DMAEMA molecule (D’ and F’). Also, the FT-IR spectrum confirms the block-copolymer PDMAEMA-b-PMPC synthesis by P-O (1240 cm^-1^) and -N(CH_3_)_2_ (3000 cm^-1^) stretching. Concerning the PDMAEMA-b-PMPC ^1^H NMR results, the peaks 2.4 (O’) and 2.8 (I’) agree with the increase of the component PDMAEMA in the two block-copolymers proposed, which corroborates with the chemical composition of the final block-copolymer. Our ^1^H NMR and FT-IR spectra results are also in concordance with previous polymer synthesis studies (Huang et al., 2017; Perecin et al., 2022; Taktak et al., 2015; Tian et al., 2020; Xie et al., 2018). The purification protocol for polymer synthesis is easy, fast, and reproducible with the total elimination of residual monomers, an essential aspect of biological applications (Jamahiriya et al., 2010), and can be applied in other studies with polymerization by RAFT.

Finally, while PMPC did not show a positive charge in any DP, our block copolymer ζ-potential results demonstrated an increase in positive charge at pH 7.4, conforming to a higher DP between PDMAEMA and PMPC. Our work demonstrated an easy and reproducible methodology to verify the block-copolymer synthesis by the ζ potential in a range of pH, which could be applied in other studies focusing on PDMAEMA and PMPC polymer approaches. Considering the above results, the polymerization of PMPC, PDMAEMA, and block-copolymer PDMAEMA-b-PMPC are clear.

Our results also demonstrated the low-toxicity polymer synthesized by different biological approaches. PMPC has been studied by previous works alone and in block copolymers, and in concordance with our results, PMPC demonstrated no significant reduction of cell viability, even with higher concentrations until 2 mg/mL. (Peng et al., 2021; Perecin et al., 2022; Zhang et al., 2022). Previous works reported the high cytotoxicity of PDMAEMA, including in concentrations lower than 200 µg/mL (Bonkovoski et al., 2014; Mendrek et al., 2018; Yu et al., 2012). Our study demonstrated the absence of cytotoxicity in the tested concentrations of PDMAEMA, potentially attributed to its lower DP. The PDMAEMA-b-PMPC block copolymers showed a slight decrease in cell viability based on metabolic activity, but no significant impact was observed using other biological toxicity measures. Despite some existing studies on the cytotoxicity of polymers synthesized by the RAFT method (Fairbanks et al., 2015; Pissuwan et al., 2010), there is an urgent need for further research to comprehensively assess their toxicity using diverse biological aspects. Remarkably, this study is the first to evaluate the cytotoxicity of PMPC, PDMAEMA, and PDMAEMA-b-PMPC block copolymers using multiple biological approaches.

The block-copolymer component proposed by our group allows future modifications by adding new blocks through RAFT polymerization. Some examples would understand the therapeutic application by delivering acid nuclei bound to PDMAEMA or changing the hydrophilic−lipophilic balance to achieve control release of hydrophilic and hydrophobic drugs.

Concerning the coating process, the IONPP demonstrated an increase in average size. Also, IONPP showed a positive surface charge after coating. The IONPP positive charge remained even after using the magnet. Moreover, the block-copolymers have a broad size distribution, with multiple populations in water solution at pH 7 (data not showed) explained by the hydrophilicity of both polymers, which cannot be detected in the IONPP sample with only one size peak. Therefore, we can infer that the cationic component of the block-copolymer is responsible for binding on the surface of the IONPs, once IONP had a negative charge, while the block-copolymer had a positive one. This coating induces colloidal stabilization, which explains the DLS results with a unique population and a good PdI (< 0.25). The coating process of IONPP by electrostriction interaction has been studied in previous works (Barrow et al., 2015; Nayeem et al., 2021). However, here we demonstrated for the first time the possibility of the coating process taking very little time and simple equipment, even with two hydrophilic polymers. This is the opposite of the standard coating process, with robust equipment (N. Zhu et al., 2018).

The FT-IR spectrum also corroborates with coating results showing the characteristic bands from the block-copolymer, the Fe-O, and sulfur groups before and then using the magnet. Also, besides the clear visualization of IONP in the bottom of the microtube (data not shown), the superparamagnetism confirms the presence and magnetic characteristics of the IONP. The M_s_ reduction is expected due to the proportion between IONP and PDMAEMA-b-PMPC, which is aligned with another coating IONP studies (Baykal et al., 2012; Perecin et al., 2022; Sood et al., 2017). We observed a more significant M_s_ reduction in our results compared to the literature. This result could be explained by the balance between IONP and block-copolymer mols. However, more studies are necessary to understand the impact of different polymer coating processes on the magnetic properties of IONP. Furthermore, TGA results demonstrated transitional behavior for IONPP ^magnet+^, confirming the stability of encapsulation. Our TEM results demonstrated IONPs coated by block-copolymer PDMAEMA-b-PMPC with an average size lower than IONPP hydrodynamic size. Therefore, it indicates some aggregation in the water solution of the IONPPs, which is also observed in other polymeric coating processes by PMPC polymer (Perecin et al., 2022).

Aptamers have been used in biomedical applications in the last few years, including for disease diagnosis. Here, we used aptamer to demonstrate one proof of concept to use the IONPP proposed by our group. First, we analyzed the binding capacity between the block-copolymer PDMAEMA-b-PMPC and HER2-5’FAM aptamer molecule. This binding occurs probably due to the interaction between aptamer phosphate and block copolymer nitrogen. The PDMAEMA-b-PMPC_10:1_ exhibited a higher aptamer binding affinity, as evidenced by its slower EC_50_ value, indicating a greater capacity for aptamer loading between the block copolymer synthesized. Due to this advantageous characteristic, this specific block copolymer was selected for further steps in the study. Therefore, our results indicated the potential binding of cationic polymer PDMAEMA concerning aptamer in a block-copolymer conformation. Cationic polymers have been used in the literature to bind to acid nuclei (Piotrowski-Daspit et al., 2020). Nevertheless, here we also calculated the EC_50_ for each polymer synthesized, including the block copolymer, making it possible to understand the impact of the cationic polymer. Also, further studies can focus on the employed polymers’ acid nuclei-carrying capacity to study the polymeric nanostructures’ transfection capacity.

The IONPPP demonstrated a greater z-average size concerning IONPP with good PdI and a slower surface charge. Moreover, we observed a clear band after using SDS to release the aptamer in agarose gel. These results emphasize the functionalization between IONPP and aptamers (IONPPP). In addition, our results applying magnet demonstrated that functionalization remains by visualization of a band in agarose gel. This way, our study shows, for the first time, the capacity of the coating and aptamer functionalized on IONP both by the electrostatic interaction.

Finally, we conducted a biological proof of concept to evaluate the binding capacity of IONPPP for aptamers, specifically targeting the HER2 protein in SKBR3 and MDA-MB-231 cells. Our results consistently demonstrated a higher number of positive cells for SKBR3 compared to MDA-MB-231, indicating the superior sensitivity achieved with the use of IONPPP. However, we observed a reduction in the efficiency of IONPPP after performing a pull-down with a magnet, in contrast to IONPPP without a magnet or the block-copolymer-aptamer complex. This decrease in efficiency could be attributed to the internalization capability of the HER2 aptamer used in this proof of concept ((K. Wang et al., 2015)). Additionally, the FAM molecule, which emits fluorescence, exhibits lower intensity in acidic pH conditions (Schneider et al., 2010). As a result, it is possible that the IONPPP was internalized by SKBR3 cells and lost its fluorescence, potentially leading to an underestimation of our results since the cells were not fixed during the assay. Moreover, after incubation with the magnet, we carefully removed all supernatant, leaving only the cell pellet captured by the magnet. We hypothesize that the binding between IONPPP and cells might not be stable, which explains the high number of negatively fluorescent cells observed when analyzing only the cells pulled by the magnet. These two hypotheses offer plausible explanations for our study’s lack of observed enrichment.

## 8. Conclusion

We present a new nanostructure called the IONPPP, a hybrid nanoparticle with an iron oxide core coated by a block-copolymer PDMAEMA-b-PMPC. Our results showed a hybrid nanoparticle with attractive chemical-physical properties for biomedical applications, such as the identification and isolation of cells by aptamers based on the presence of surface proteins.

## Abbreviations

(DLS): Dynamic Light Scattering
(PdI): Polydispersity Index
(TGA): Thermogravimetry Analysis
(PDMAEMA): Poly(2-(Dimethylamino)Ethyl Methacrylate)
(PMPC): Poly (2-Methacryloyloxyethyl Phosphorylcholine)
(IONPs): Iron Oxide Nanoparticles
(NP): Nanoparticle
(RAFT): Reversible Addition−Fragmentation Chain Transfer
(XRD): X-Ray Diffractometry
(M_s_): Magnetization Value
(TEM): Transmission Electron Microscopy
(HRTEM): High-Resolution Transmission Electron Microscopy
(FT-IR): Fourier-transform infrared spectroscopy
(^1^H NMR): ^1^H Nuclear Magnetic Resonance Spectroscopy
(SAED): Selected Area Electron Diffraction
(VSM): Vibrating Sample Magnetometer
(HER2): human epidermal growth factor receptor 2
(SELEX): Systematic Evolution of Ligands by Exponential Enrichment
(M_n_): number average molar mass
(DP): degree of polymerization
MTT: 3-(4,5 Dimethyl-2-thiazolyl)-2,5-diphenyl-2H-tetrazolium Bromide
(PI): propidium iodide
(DAPI): 4’, 6-Diamidino-2-Phenylindole
(SD): standard deviation

## Acknowledgments

The authors would like to acknowledge financial support from the Brazilian agency Coordenação de Aperfeiçoamento de Pessoal de Nível Superior (CAPES) and São Paulo Research Foundation (FAPESP) for the awards #19/24563-0 (ASJJ), and #19/16351-3 (SMG). The LNNano to microscopy, XRD analysis of nanoparticles, and the LQS at LNBio all help with polymer synthesis. Also, we thank LNBio and Bionanomanufacturing Center for access to core facilities and financial support. We would like to thank the Fundação de Apoio ao Instituto de Pesquisas Tecnologicas (FIPT) for support from the Young Talent program.

