## Supplementary information for "Iron Oxide Nanoparticles Coated with Biodegradable Block-Copolymer PDMAEMA-b-PMPC and Functionalized with Aptamer for HER2 Breast Cancer Cell Identification"

Dr. Natália Neto Pereira Cerize

Zip code: 05508-070

Bionanomanufacturing Center, Institute for Technological Research (IPT)

Av. Prof. Almeida Prado, 1032 - Butantã, São Paulo – SP – Brazil

*** Corresponding author**

Dr. Sandra Dias;

Zip code: 13083-100

+55 19 3512-3503

Brazilian Biosciences National Laboratory (LNBio), Brazilian Center for Research in Energy and Materials (CNPEM)

Rua Giuseppe Máximo Scolfaro, 10.000 - Polo II de Alta Tecnologia de Campinas – SP

### Supplementary information

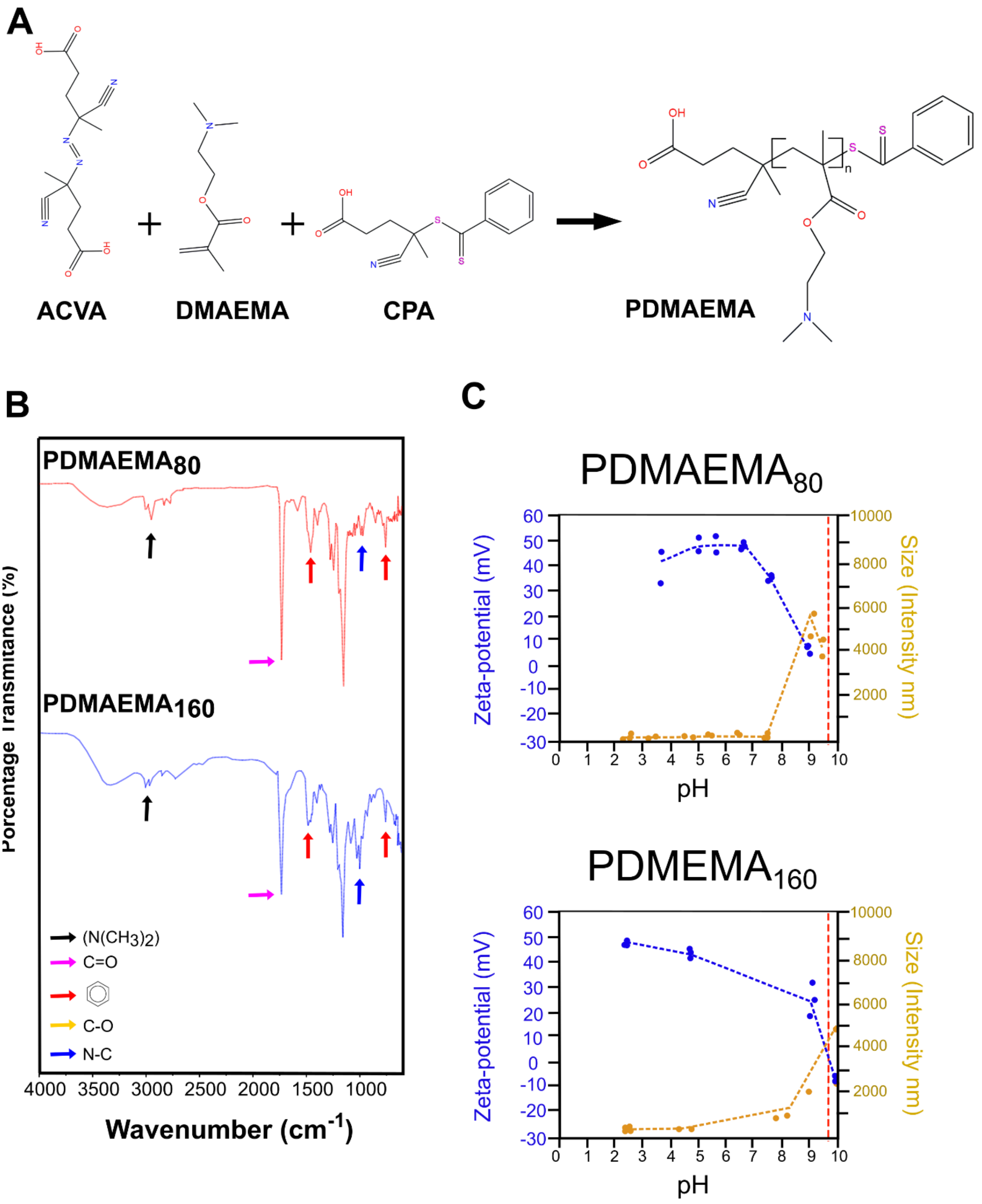


**Figure Supplementary 1. Polymer characterization. A)** Scheme representation of experimental approach synthesis of the hydrophilic cationic PDMAEMA; **B)** FT-IR (Fourier-transformation infrared spectroscopy) spectrum PDMAEMA results; **C)** z-average size and ζ-Potential of PDMAEMAs in different pHs by DLS (Dynamic light scattering).

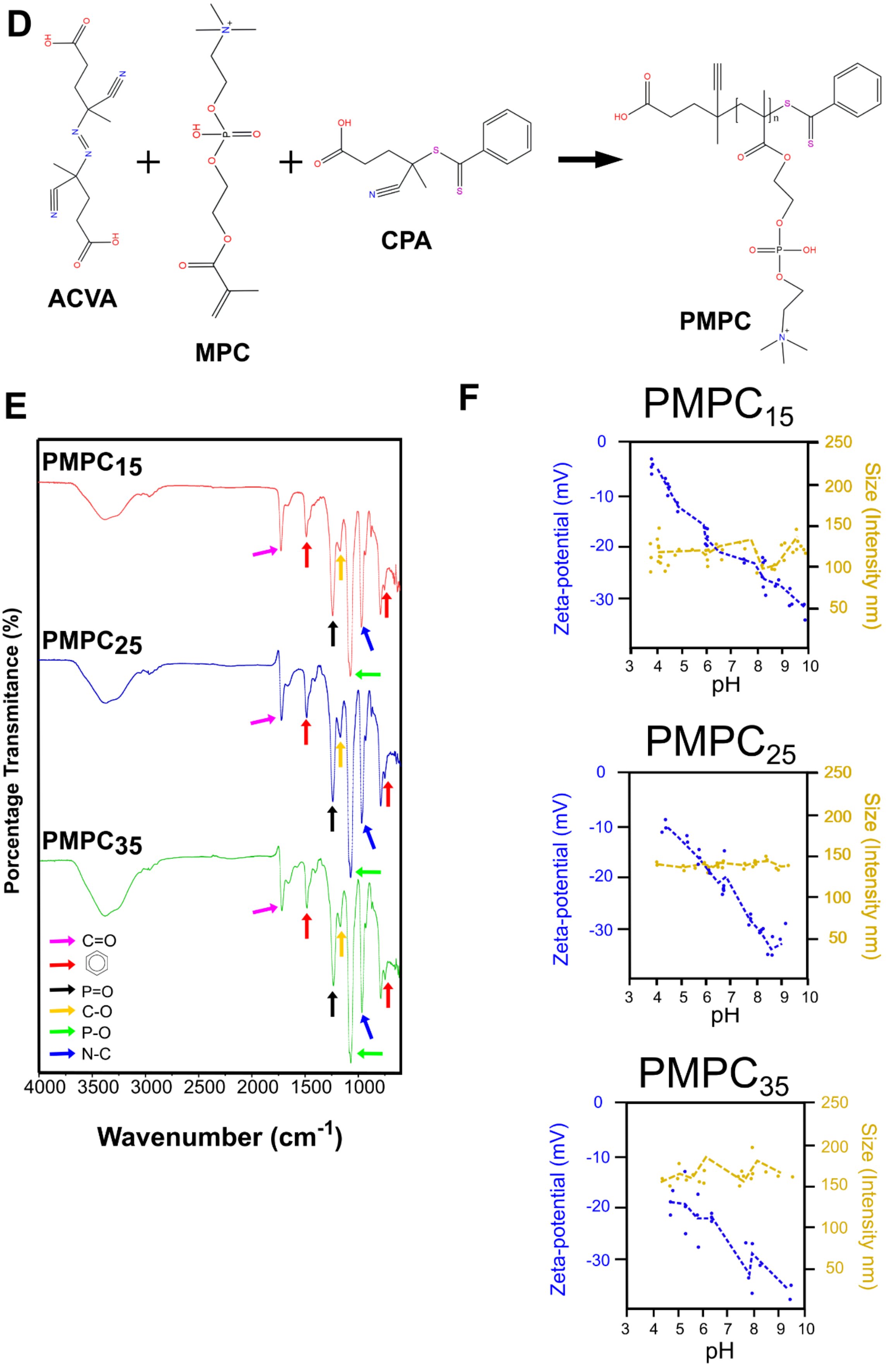


**Figure Supplementary 1. Polymer characterization. D)** Scheme representation of experimental approach synthesis of the zwitterionic polymer PMPC; **E)** FT-IR (Fourier-transformation infrared spectroscopy) spectrum PMPC results; **F)** z-average size and ζ-Potential of PMPCs in different pHs by DLS (Dynamic light scattering).

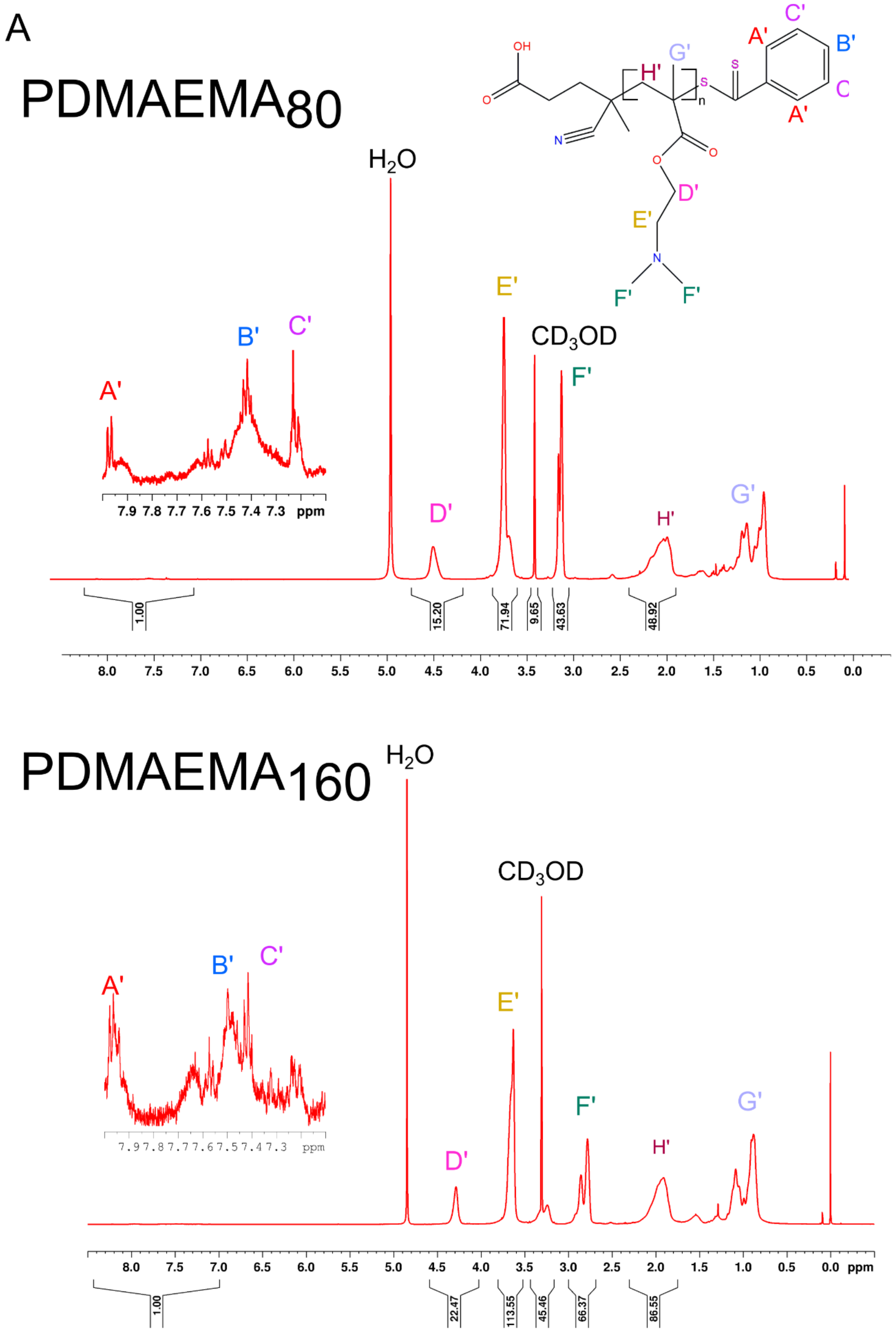


**Figure Supplementary 2. [1]H NMR spectra results. A)** [1]H NMR spectra results for PDMAEMAs,

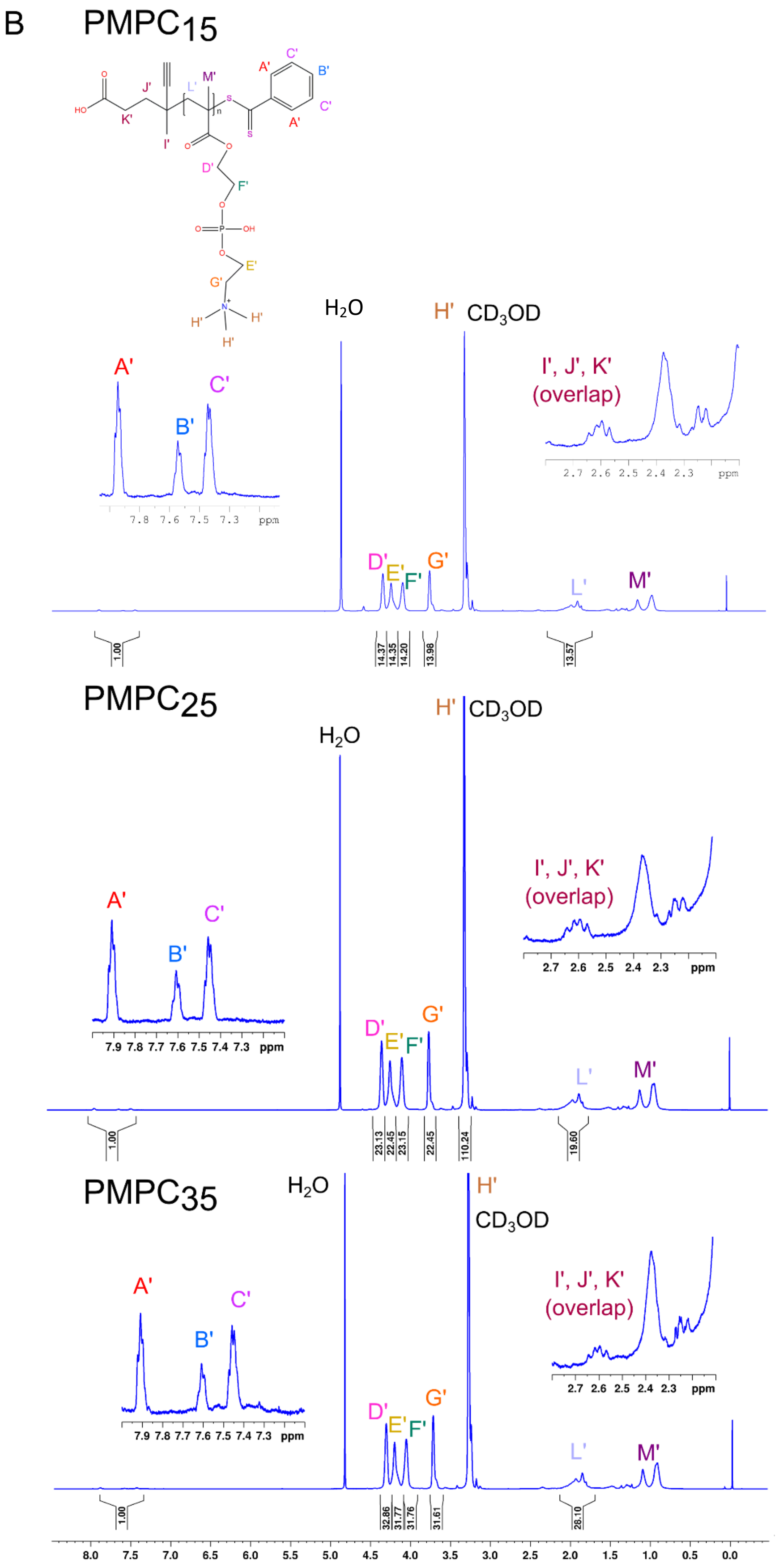


**Figure Supplementary 2. [1]H NMR spectra results. B)** [1]H NMR spectra results for PMPCs.

C

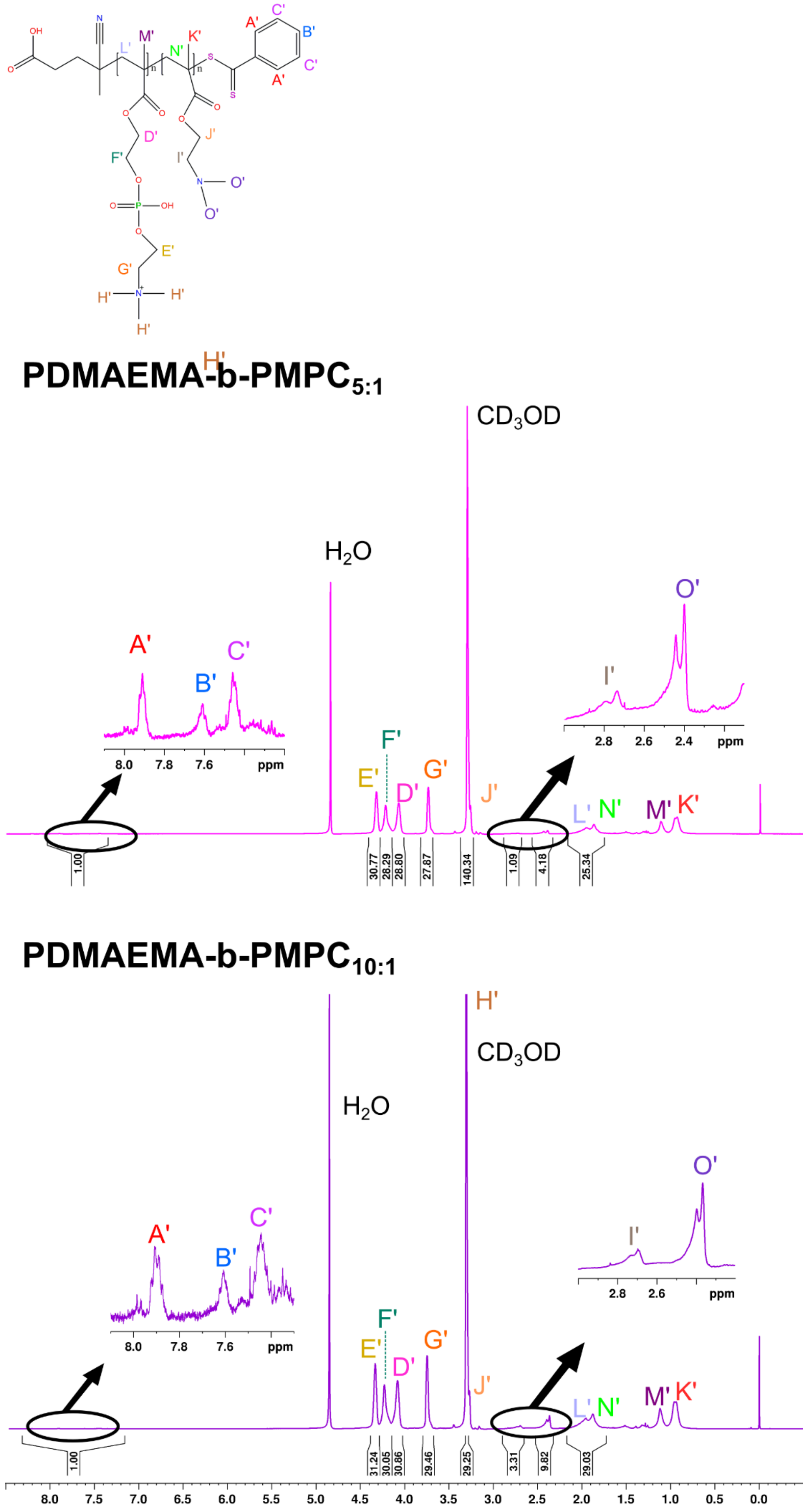


**Figure Supplementary 2. $^{1}$H NMR spectra results. C)** $^{1}$H NMR spectra results for block-copolymers PDMAEMA-b-PMPCs.

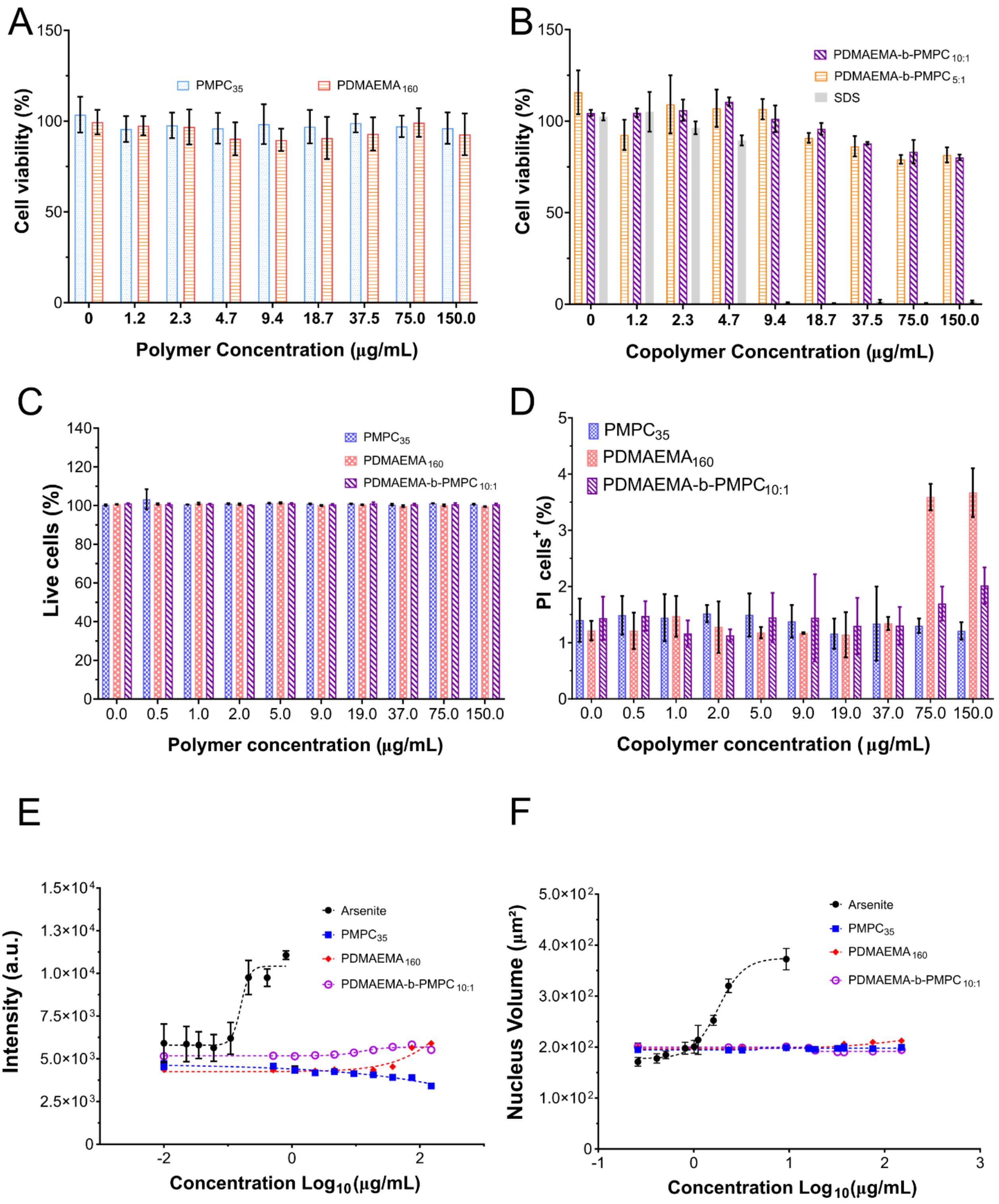


**Figure Supplementary 3. Cytotoxicity panel.** The MTT assay for **A)** PDMAEMA and PMPC polymer and **B)** block-copolymers PDMAEMA-b-PMPC, **C)** the percentage of live cells, and **D)** PI% relative cell staining in HaCaT cells incubated with polymers (PDMAEMA and PMPC) and block-copolymer (PDMAEMA-b-$PMPC_{10:1}$); **E)** mitochondria weight and **F)** nucleus volume of HaCaT cells incubated with polymers (PDMAEMA and PMPC) and block-copolymer (PDMAEMA-b-$PMPC_{10:1}$) in comparison with death molecule control (arsenite).

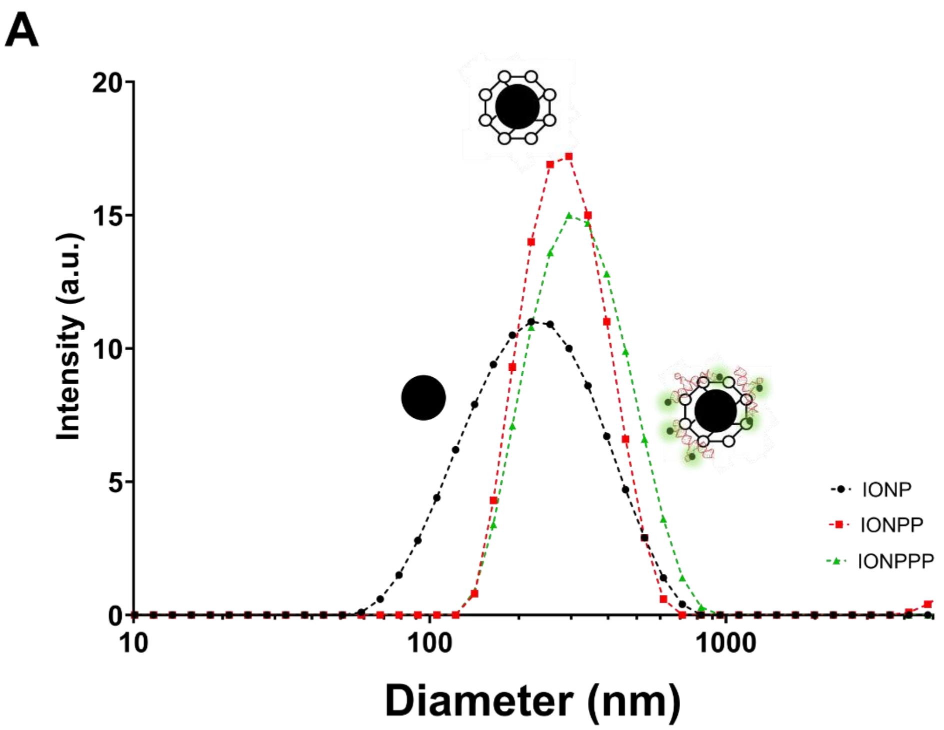


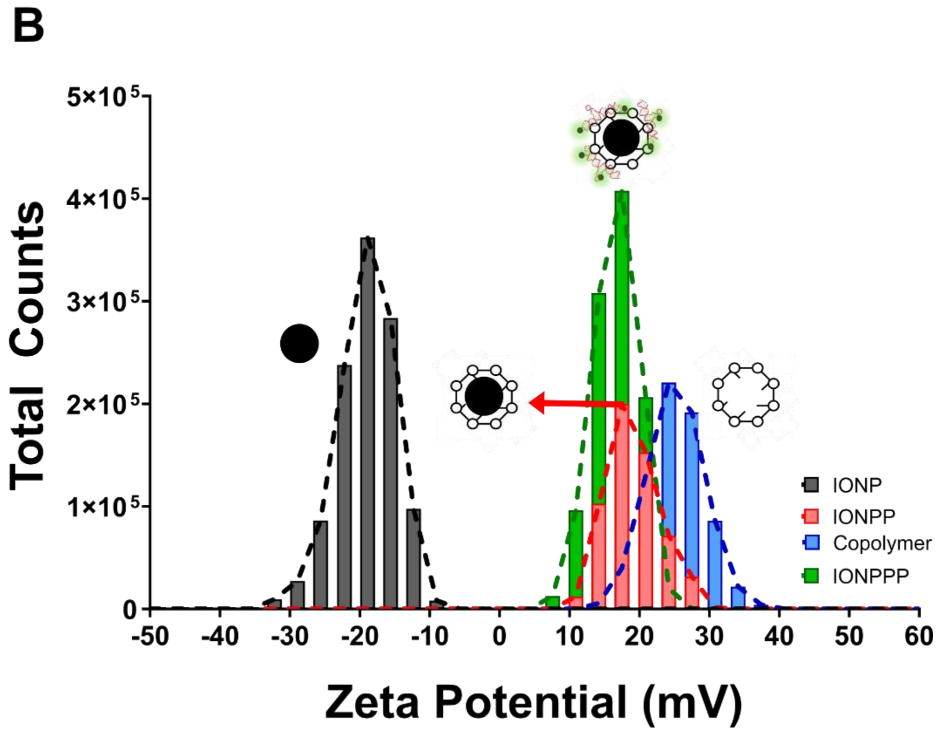


**Figure Supplementary 4. IONPPP characterization. A)** z-average size reported by the intensity of IONP, IONPP, block-copolymer (PDMAEMA-b-PMPC), and IONPP by DLS, and **B)** ζ-Potential of IONP, IONPP, block-copolymer (PDMAEMA-b-PMPC), and IONPP by DLS (Dynamic light scattering).

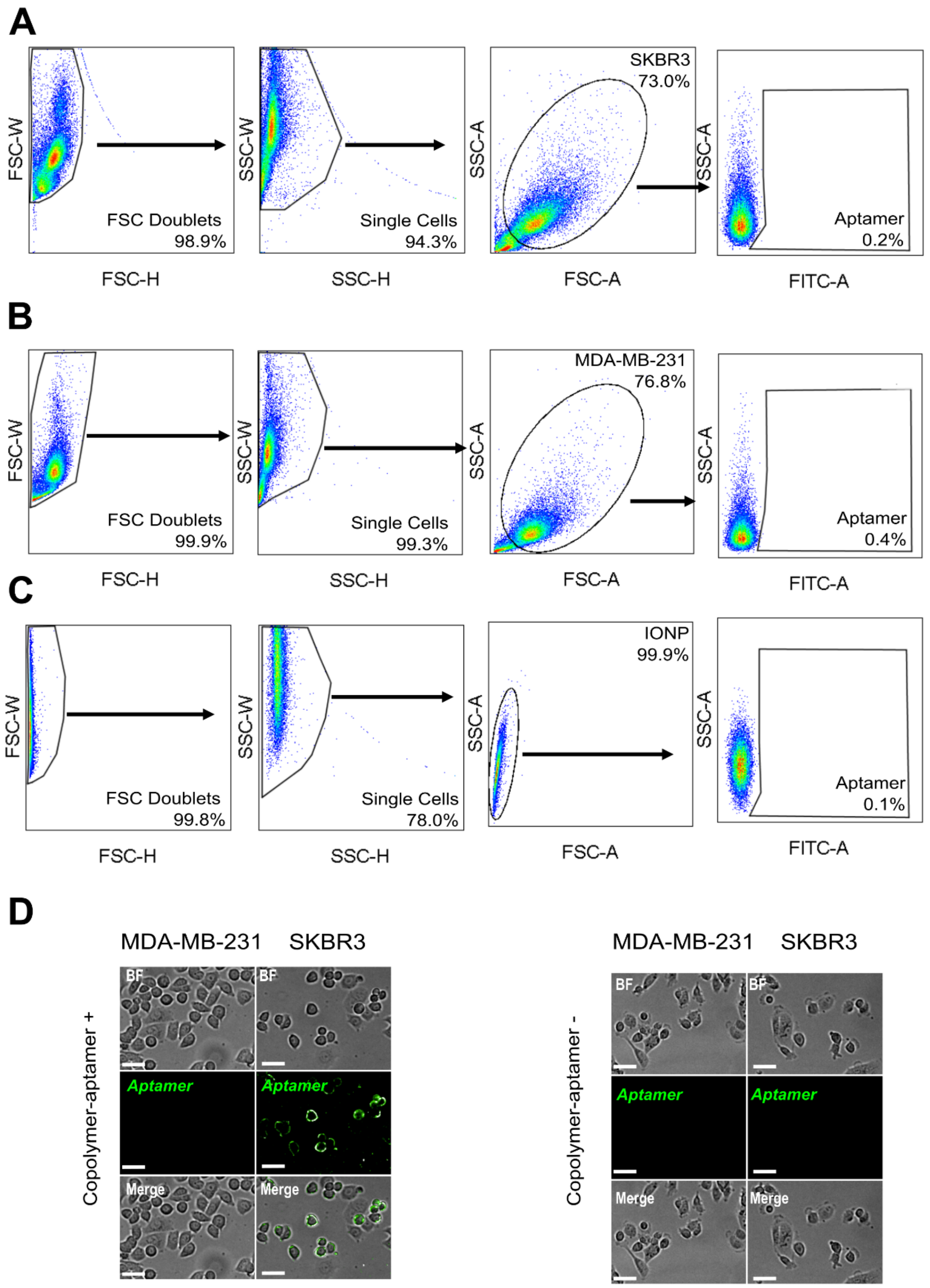


**Figure Supplementary 5. Gate strategy and microscopy analysis.** Flow cytometry results related to gate strategy for **A)** SKBR3 and **B)** MDA-MB-231 tumor cells; **C)** IONP profile results in flow cytometry; **D)** fluorescence microscopic images of the PDMMA-b-PMPC – aptamer complex incubated in HER-2 cell$^{-/+}$.
